# Elementary Integrate-and-Fire Process Underlies Pulse Amplitudes in Electrodermal Activity

**DOI:** 10.1101/2021.01.06.425499

**Authors:** Sandya Subramanian, Patrick L. Purdon, Riccardo Barbieri, Emery N. Brown

## Abstract

Electrodermal activity (EDA) is a direct read-out of sweat-induced changes in the skin’s electrical conductance. Sympathetically-mediated pulsatile changes in skin sweat measured as EDA resemble an integrate-and-fire process, which yields an inverse Gaussian model as the inter-pulse interval distribution. We have previously showed that the inter-pulse intervals in EDA follow an inverse Gaussian distribution. However, the statistical structure of EDA pulse amplitudes has not yet been characterized based on the physiology. Expanding upon the integrate-and-fire nature of sweat glands, we hypothesized that the amplitude of an EDA pulse is proportional to the excess volume of sweat produced compared to what is required to just reach the surface of the skin. We modeled this as the difference of two inverse Gaussian models for each pulse, one which represents the time required to produce just enough sweat to rise to the surface of the skin and one which represents the time requires to produce the actual volume of sweat. We proposed and tested a series of four simplifications of our hypothesis, ranging from a single difference of inverse Gaussians to a single simple inverse Gaussian. We also tested four additional models for comparison, including the lognormal and gamma distributions. All models were tested on EDA data from two subject cohorts, 11 healthy volunteers during 1 hour of quiet wakefulness and a different set of 11 healthy volunteers during approximately 3 hours of controlled propofol sedation. All four models represent simplifications of our hypothesis outperformed other models across all 22 subjects, as measured by Akaike’s Information Criterion (AIC), as well as mean and maximum distance from the diagonal on a quantile-quantile plot. Our broader model set of four simplifications offered a useful framework to enhance further statistical descriptions of EDA pulse amplitudes. Some of the simplifications prioritize fit near the mode of the distribution, while others prioritize fit near the tail. With this new insight, we can summarize the physiologically-relevant amplitude information in EDA with at most four parameters. Our findings establish that physiologically based probability models provide parsimonious and accurate description of temporal and amplitude characteristics in EDA.

**AUTHOR SUMMARY:** Electrodermal activity (EDA) is an indirect read-out of the body’s sympathetic nervous system, or fight-or-flight response, measured as sweat-induced changes in the electrical conductance properties of the skin. Interest is growing in using EDA to track physiological conditions such as stress levels, sleep quality, and emotional states. Our previous worked showed that the times in between EDA pulses obeyed a specific statistical distribution, the inverse Gaussian, that arises from the physiology of EDA production. In this work, we build on that insight to analyze the amplitudes of EDA pulses. In an analysis of EDA data recorded in 11 healthy volunteers during quiet wakefulness and 11 different healthy volunteers during controlled propofol sedation, we establish that the amplitudes of EDA pulses also have specific statistical structure, as the differences of inverse Gaussians, that arises from the physiology of sweat production. We capture that structure using a series of progressively simpler models that each prioritize different parts of the pulse amplitude distribution. Our findings show a physiologically-based statistical model provides a parsimonious and accurate description EDA. This enables increased reliability and robustness in analyzing EDA data collected under any circumstance.

## INTRODUCTION

Sweat gland activity is used to assess sympathetic nervous system activity in applications such as lie detector tests and neuromarketing (1). Sympathetic activation is also known as the “fight or flight response”, which is induced by states such as stress, anxiety, and pain (1). Electrodermal activity (EDA) measures the second-to-second electrical conductance of the skin to capture sweat gland activity. As stimulation of sweat glands increases due to stress or pain for example, more sweat is produced, which increases the electrical conductance of the skin. EDA is typically divided into two components (1). The first is a baseline or tonic component which drifts gradually over minutes and is thought to represent ambient conditions which contribute to baseline level of filling of the glands. The second is the phasic component, which rides on top of the tonic and consists of pulsatile sweat release events. These pulsatile sweat release events have a timescale of a few seconds and are thought to correspond more closely to sympathetic nervous system activity (1). There is growing interest in the development of algorithms to accurately characterize changes in emotional and physiologic states from EDA.

Our previous analyses have showed that the inter-pulse interval distribution in EDA data follows an inverse Gaussian distribution, which agrees with a model of the rise of sweat through the gland to the skin surface as an integrate-and-fire process, specifically a Gaussian random walk with drift diffusion (2,3). We showed that deviations from the inverse Gaussian due to recording across many sweat glands tend toward right-skewed heavier tailed distributions, such as the lognormal (2). Using these insights, we further defined a low-order paradigm for verifying the physiologic structure in EDA that includes a framework for goodness-of-fit analysis (4,5).

However, temporal information is not the only information in phasic EDA. Each pulse occurs not only at a discrete point in time, but also with a specific amplitude (1). Existing algorithms for phasic EDA analysis assume that the amplitude of each pulse has a one-to-one association with the intensity of the stimulus driving it (6-10); however, previous physiologic experiments have shown that the background level of nerve activity and baseline level of filling of the sweat glands can alter pulse amplitude even in the face of an unchanging stimulus intensity (11-13). In this work, we propose a model for pulse amplitudes that uses the same insight about integrate-and-fire physiology of sweat glands as for temporal information. We hypothesize that the amount of sweat produced in a pulse relates directly to stimulus amplitude. However, we postulate that the observed amplitude of the pulse relates to how much more sweat is produced than what is required to reach the surface of the skin, which also accounts for the role played by the background filling level of the sweat glands. Therefore, we model the amplitude of a pulse as the difference of actual amount of sweat produced and the baseline filling level.

We implement four different simplifications of our model in two different subject cohorts. The four simplifications tested were a simple inverse Gaussian, a three-parameter inverse Gaussian with the third parameter serving as a location shift, a single difference of an inverse Gaussian and a Gaussian, and a single difference of two inverse Gaussians. We show, using a goodness-of-fit analysis, that each simplification balances the important characteristics of the model differently. The simple inverse Gaussian model fits the mode of the pulse amplitude distribution well, while the difference models better capture the tail. The three-parameter model seems to balance both.

Important advances we report are a set of low-order physiology-based point process models for pulse amplitudes in phasic EDA that work synergistically with our existing models for temporal information. Using both together, we can extract all relevant information from phasic EDA in statistically rigorous way. The balance of this paper is organized as follows. In **Physiology & Models**, we derive our hypothesis about pulse amplitudes from the integrate-and-fire physiology, outline four statistical models to capture it that involve the inverse Gaussian as well as four alternatives for comparison. In **Application** and **Results**, we use these models in the analysis of EDA pulse amplitudes recorded from 22 subjects across two different subject cohorts, one while awake and at rest and the other under controlled propofol sedation. The **Discussion** describes the implications of our findings for future basic science and translational studies.

## MATERIALS & METHODS

### Anatomy and Physiology

To develop our statistical models of EDA pulse amplitudes, we review the anatomy and physiology of sweat production in the skin. An eccrine sweat gland has three parts: the dermal gland which secretes sweat, the duct which connects the gland to the surface of the skin, and the pore which opens the duct to the skin surface (14). The dermal gland is innervated by a peripheral sympathetic nerve called the sudomotor nerve. Sympathetically-induced increases in spiking activity in the sudomotor nerves, called sudomotor bursts, cause sweat production in the gland. Sweat produced in response to these bursts accumulates in the duct, rising up to the skin surface by pushing open the pore. At the same time, sweat dissipates by constant reabsorption through the walls of the duct and by evaporation from the skin surface (14).

Sweat increases the skin’s electrical conductance because the salt-containing sweat in the gland creates a low-resistance path through the skin, especially once it has crossed the high-resistance top layer of skin, the stratum corneum (14). The greater the number of sweat glands bursting simultaneously, the greater the increase in conductance of the skin. The electrical conductance across the skin can be measured by placing two electrodes on the palmar surface of the hand, applying a constant voltage, and measuring the current (1). The pulsatile changes in conductance measured in the skin are termed galvanic skin responses (GSRs), referred to as ‘pulses’ in this paper.

Existing algorithms for EDA analysis typically assume that pulse amplitude can be explained solely by the intensity of the stimulus, and therefore they rely on the pulse amplitude to directly infer stimulus amplitude (6-10). However, physiologic experiments done by Wallin et al. in the 1990s demonstrated that modulating the background alone could result in pulses of varying amplitudes, even if stimulus intensity was held constant (11-13). Therefore, interpreting pulse amplitude in light of stimulus intensity alone, especially in a context in which background cannot be held constant, can be misleading. Any physiologically viable model for pulse amplitude must also account for the background.

Building upon the integrate-and-fire model we postulated for temporal information in sweat gland bursts, we hypothesized that the amplitude of a pulse is proportional to the excess volume of sweat produced compared to what would be required to reach the surface of the skin, where the total volume of sweat produced relates to the intensity of the stimulus, and the amount of sweat required to reach the surface of the skin is determined by the background filling level. The background filling level is affected by tonic EDA, spontaneous activity, reabsorption rate in the duct of the sweat gland, each individual’s autonomic ‘excitability’, and the conductance properties of their skin (11-16). It also varies from sweat gland to sweat gland.

### Statistical Model

By assuming a relatively linear relationship between the volume of sweat in the sweat glands and measured electrical conductance across the skin, and a relatively constant rate of sweat production once stimulated, we can relate measured electrical conductance across the skin to the times taken to secrete the required volume of sweat. Both assumptions are simplifications to the true microfluidic properties, made across the aggregate of hundreds of sweat glands (14). With the help of these assumptions, we hypothesize that the amplitude of each pulse can be modeled as the difference of two processes: one integrate-and-fire process to reach the surface of the skin (the minimum amount of sweat production required), and a second integrate-and-fire process to reach an imaginary threshold higher than the surface of the skin (the actual sweat production resulting from the intensity of the stimulus). However, each pulse may have a different stimulus intensity and background filling level, so the two processes are not identical across pulses. However, since they are both integrate-and-fire processes, we hypothesize that each pulse amplitude can be modeled as the difference of two inverse Gaussian distributions (Fig 1A; 17-19).

**Fig 1.**
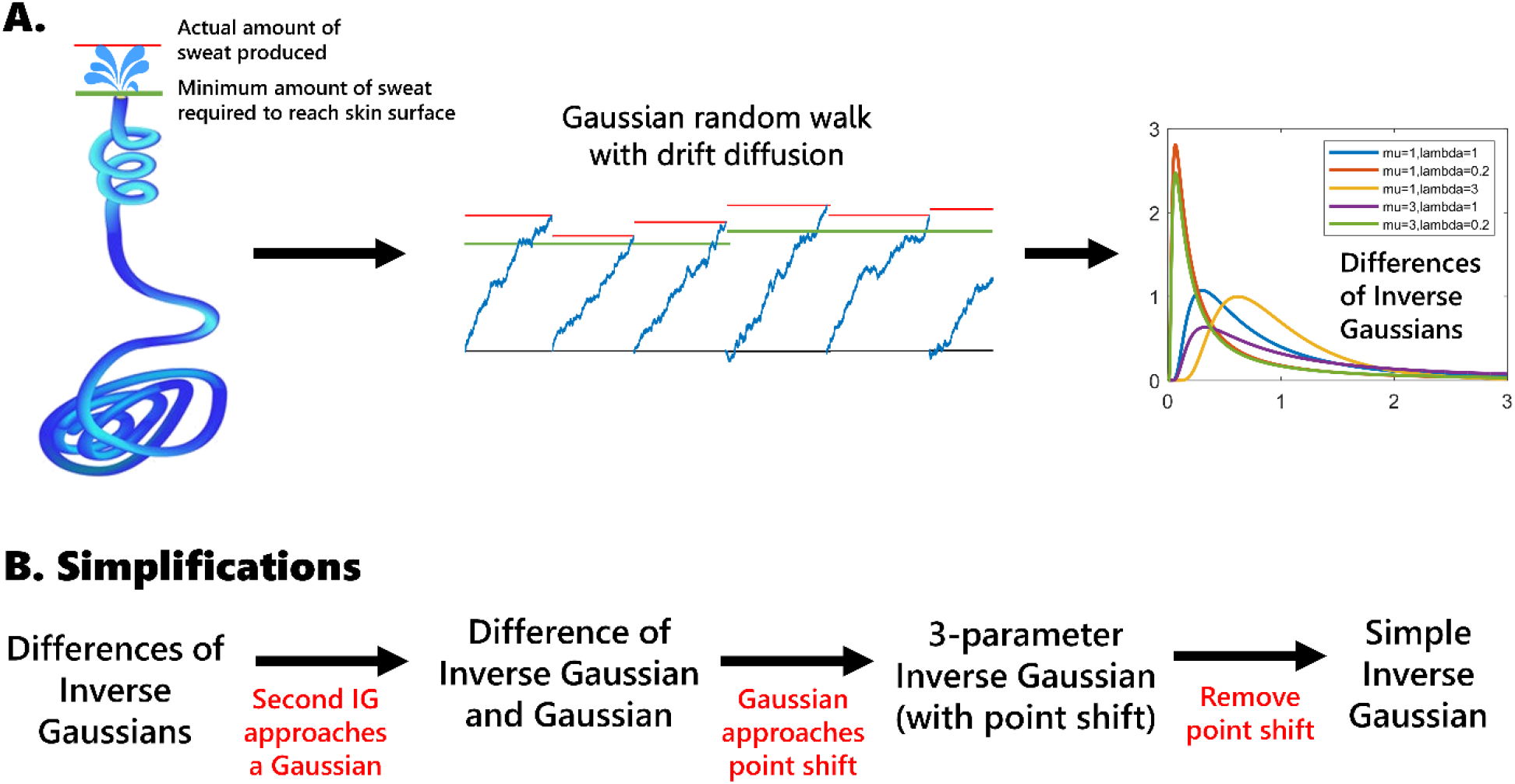
Schematic of our physiological hypothesis and statistical model simplifications. (A) We hypothesize that the amplitude of a pulse is related to the excess volume of sweat produced relative to what is required to reach the surface of the skin, which translates to the difference of first passage times between two integrate-and-fire processes, yielding the difference of two inverse Gaussian distributions.(B) We arrived at a series of simplifications of our model based on the statistical properties observed. IG = inverse Gaussian

Since fitting this model for each pulse individually is an under-constrained problem, we proposed four viable simplifications across a single subject’s dataset (Fig 1B):

1. a single difference of inverse Gaussians (IG-IG),
2. a single difference between an inverse Gaussian and a Gaussian (IG-G),
3. a single 3-parameter inverse Gaussian with the third parameter being a location shift (3IG), and
4. a single simple inverse Gaussian model (SIG).

The first simplification is the most obvious place to start, but preliminary results indicated that the second inverse Gaussian actually approached a very narrow Gaussian distribution (19), leading to the second and third simplifications. The fourth simplification is the simplest of all four. Our previous work indicated that the inverse Gaussian was the best integrate-and-fire model for EDA data across subjects (2), and therefore we only included the inverse Gaussian rather than the larger family of all integrate-and-fire models. We also compared other families of right-skewed models, such as the lognormal, exponential, and gamma. We performed a goodness-of-fit analysis for all models with quantile-quantile (QQ) plots and rescaled QQ plots. QQ plots show the quantiles of one distribution against another, in this case comparing the theoretical and empirical distributions (20,21). We also calculated Akaike’s Information Criterion (AIC) and the mean and maximum distance from the 45-degree line on the QQ plots (20,21).

### Experimental Data

We tested all of the models on two subject cohorts, collected at different times using different equipment and under different conditions. The first cohort of data is EDA data we previously collected from 12 healthy volunteers between the ages of 22 and 34 (6 males) while awake and at rest (2). The study was approved by the Massachusetts Institute of Technology (MIT) Committee on the Use of Humans as Experimental Subjects. All subjects provided written informed consent. Approximately one hour of EDA data was collected at 256 Hz from electrodes connected to the second and fourth digits of each subject’s non-dominant hand. Subjects were seated upright and instructed to remain awake. They were allowed to read, meditate, or watch something on a laptop or tablet, but not to write with the instrumented hand. One subject’s data were not included in the analysis because we learned after completing the experiment that the subject occasionally experienced a Raynaud’s type phenomenon, which would affect the quality of the EDA data. Data from the remaining 11 subjects were analyzed.

The second data cohort consists of EDA recorded from eleven healthy volunteers during a study of propofol-induced unconsciousness (22). The protocol was approved by the Massachusetts General Hospital (MGH) Human Research Committee. All subjects provided written informed consent. For all subjects, approximately 3 hours of data were recorded while using target-controlled infusion protocol. The data collection is described in detail in (22). The infusion rate was increased and then decreased in a total of ten stages of roughly equal lengths to achieve target propofol concentrations of: 0 mg/kg/hr, 1, 2, 3, 4, 5, 3.75, 2.5, 1.25, 0. All data were analyzed using Matlab R2019a.

### Data Availability Statement

All data files will be available from the PhysioNet database. The awake and at rest cohort data is already available at (23).

### Data Preprocessing and Pulse Selection

Preprocessing consisted of two major steps, 1) detecting and removing artifacts and 2) isolating the phasic component. Both have been described in previously in (5). Because of the level of high frequency noise seen in the recording equipment used for the propofol data, those data were additionally low-pass filtered with a cutoff of 3 Hz after artifact removal.

Pulse selection was done using the methodology described in (4) in which the fits of four right-skewed models were used to select the best prominence threshold at which to extract pulses. Prominence is a locally adjusted amplitude measure computed using the *findpeaks* algorithm in Matlab. This algorithm adjusts the amplitude of each peak in the signal as the height above the highest of neighboring “valleys” on either side. The valleys are chosen based on the lowest point in the signal between the peak and the next intersection with the signal of equal height on either side. Since the same pulse selection framework was followed on the same two cohorts of data, the pulses selected for each subject were also the same as in (4). The final thresholds used for each subject and temporal properties of the pulses selected can be found in (4). For this paper, we used the prominence of the extracted pulses as the measure of pulse amplitude.

### Statistical Model Fitting and Comparison

We fitted eight models to each subject’s dataset of extracted pulse amplitudes (prominences). The first four were the four simplifications of our hypothesis, and the other four were other models for comparison. The eight models fitted were:

1. a single difference of inverse Gaussians (IG-IG),
2. a single difference between an inverse Gaussian and a Gaussian (IG-G),
3. a single 3-parameter inverse Gaussian with the third parameter being a fitted location shift (3IG with fitted shift),
4. a single simple inverse Gaussian model (SIG),
5. a single lognormal model (L),
6. a single gamma model (G),
7. a single exponential model (E), and
8. a single 3-parameter inverse Gaussian with the location shift parameter set to the prominence threshold used to extract pulses (3IG with known shift).

The closed form densities for Models 3-8 are in S2 Appendix. We fitted models 4-7 by maximum likelihood (24). We fitted models 1-3 and 8 by method-of-moments (25), due to a lack of closed form solutions for maximum likelihood estimates of the parameters. The derivation of method of moments estimates for Models 1 and 2 is detailed in S3 Appendix. Method of moments estimates for Model 3 (also used for Model 8) were given in (19). Model 8 was included to verify that the process of pulse extraction using a prominence threshold did not skew the pulse amplitude results. We assessed goodness-of-fit by Akaike’s Information Criterion (AIC) and QQ plots. The AIC is defined as

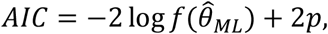

where 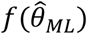 is the likelihood evaluated at the maximum likelihood estimate of the parameters and is the number of parameters. A lower AIC indicates a better fit. For the models fitted by method of moments, we estimated the log likelihood numerically. AIC prioritizes efficiency of the model.

We also plotted QQ plots and rescaled QQ plots. For rescaled QQ plots, both model and empirical quantile values were rescaled by

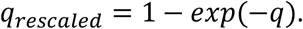

Rescaled QQ plots were used for more uniform visualization of the data across all quantiles. From the QQ plots (not rescaled), we calculated the mean and maximum perpendicular distances between the plotted fit and the 45-degree line, which represents a perfect fit. A lower mean distance indicates a better average fit across all quantiles, while a lower maximum distance indicates a better worst case fit (a better worst-fitting point).

## RESULTS

### Extraction of pulses

In the awake and at rest cohort, the number of pulses extracted per subject across one hour ranged from 97 to 324 using prominence thresholds ranging from 0.0025 to 0.027. In the propofol sedation cohort, the number of pulses extracted per subject across 3-4 hours ranged from 383 to 1250 using prominence thresholds ranging from 0.02 to 0.055.

### Findings from statistical model comparison

Based on AIC, in the awake and at rest cohort (Tables 1-2), SIG is the best model for 9 out of the 11 subjects (S3-S11), lognormal for one subject (S2), and the 3IG with known location shift (Model 8) for one subject (S1). The SIG was always the best of the four simplifications (Models 1-4). In the propofol sedation cohort (Tables 3-4), SIG was the best model for 8 of 11 subjects (P1-P7, P11), 3IG with fitted shift for one (P8) and lognormal for two (P9, P10).

**Table 1.**
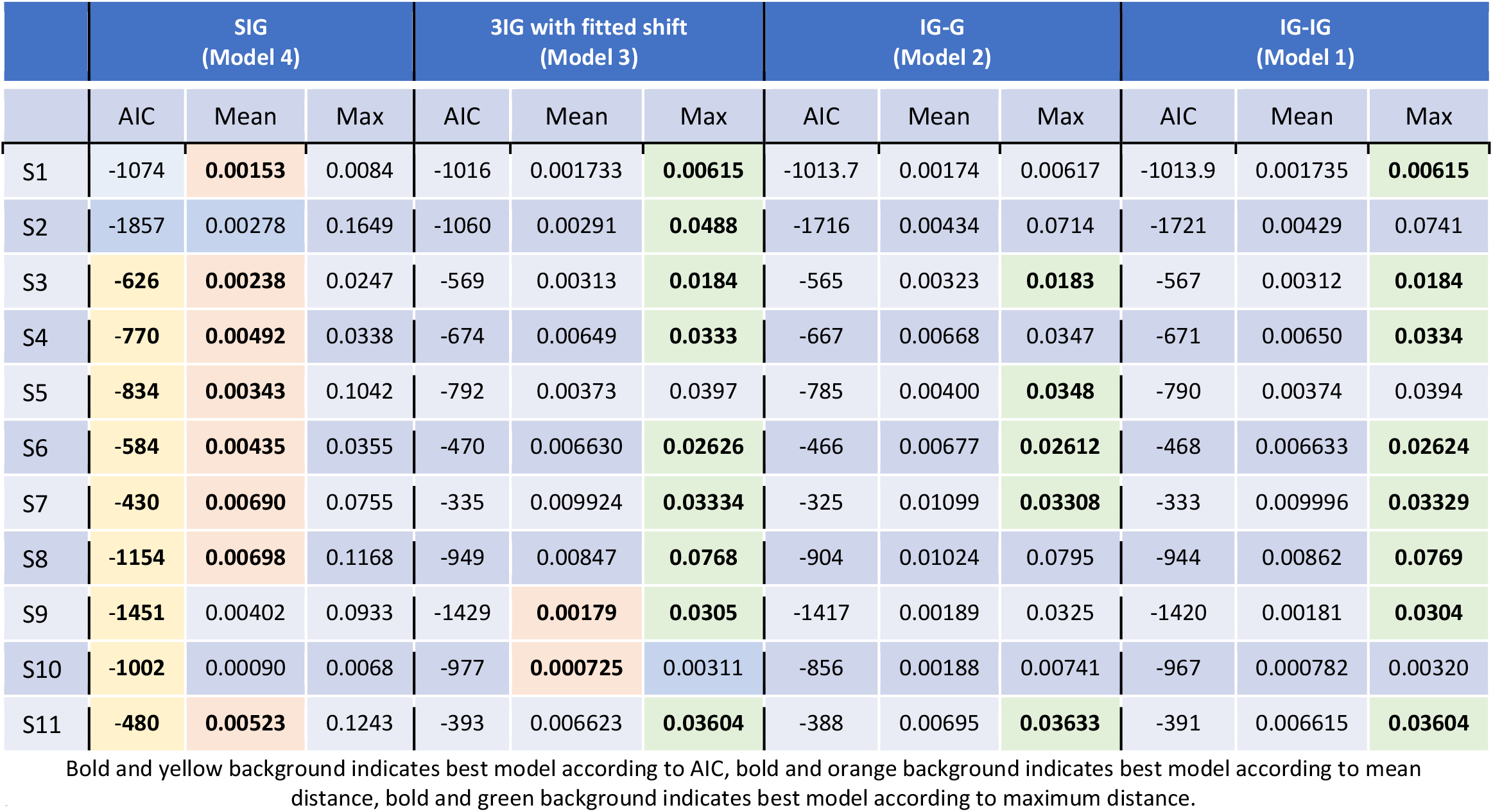
Model fit results for awake and at rest cohort for Models 1-4.

|  | SIG<br>(Model 4) |  |  | 3IG with fitted shift<br>(Model 3) |  |  | IG-G<br>(Model 2) |  |  | IG-IG<br>(Model 1) |  |  |
| --- | --- | --- | --- | --- | --- | --- | --- | --- | --- | --- | --- | --- |
|  | AIC | Mean | Max | AIC | Mean | Max | AIC | Mean | Max | AIC | Mean | Max |
| S1 | -1074 | **0.00153** | 0.0084 | -1016 | 0.001733 | **0.00615** | -1013.7 | 0.00174 | 0.00617 | -1013.9 | 0.001735 | **0.00615** |
| S2 | -1857 | 0.00278 | 0.1649 | -1060 | 0.00291 | **0.0488** | -1716 | 0.00434 | 0.0714 | -1721 | 0.00429 | 0.0741 |
| S3 | **-626** | **0.00238** | 0.0247 | -569 | 0.00313 | **0.0184** | -565 | 0.00323 | **0.0183** | -567 | 0.00312 | **0.0184** |
| S4 | **-770** | **0.00492** | 0.0338 | -674 | 0.00649 | **0.0333** | -667 | 0.00668 | 0.0347 | -671 | 0.00650 | **0.0334** |
| S5 | **-834** | **0.00343** | 0.1042 | -792 | 0.00373 | 0.0397 | -785 | 0.00400 | **0.0348** | -790 | 0.00374 | 0.0394 |
| S6 | **-584** | **0.00435** | 0.0355 | -470 | 0.006630 | **0.02626** | -466 | 0.00677 | **0.02612** | -468 | 0.006633 | **0.02624** |
| S7 | **-430** | **0.00690** | 0.0755 | -335 | 0.009924 | **0.03334** | -325 | 0.01099 | **0.03308** | -333 | 0.009996 | **0.03329** |
| S8 | **-1154** | **0.00698** | 0.1168 | -949 | 0.00847 | **0.0768** | -904 | 0.01024 | 0.0795 | -944 | 0.00862 | **0.0769** |
| S9 | **-1451** | 0.00402 | 0.0933 | -1429 | **0.00179** | **0.0305** | -1417 | 0.00189 | 0.0325 | -1420 | 0.00181 | **0.0304** |
| S10 | **-1002** | 0.00090 | 0.0068 | -977 | **0.000725** | 0.00311 | -856 | 0.00188 | 0.00741 | -967 | 0.000782 | 0.00320 |
| S11 | **-480** | **0.00523** | 0.1243 | -393 | 0.006623 | **0.03604** | -388 | 0.00695 | **0.03633** | -391 | 0.006615 | **0.03604** |

**Table 2.** Model fit results for awake and at rest cohort for Models 1-4.

|  | 3IG with known shift<br>(Model 8) |  |  | Exponential<br>(Model 7) |  |  | Gamma<br>(Model 6) |  |  | Lognormal<br>(Model 5) |  |  |
| --- | --- | --- | --- | --- | --- | --- | --- | --- | --- | --- | --- | --- |
|  | AIC | Mean | Max | AIC | Mean | Max | AIC | Mean | Max | AIC | Mean | Max |
| S1 | **-1098** | 0.0024 | 0.0780 | -1001 | 0.00160 | 0.0099 | -1029 | 0.00204 | 0.0101 | -1066 | 0.00162 | 0.0099 |
| S2 | -1073 | 0.0094 | 0.7340 | -1729 | 0.00362 | 0.1721 | -1747 | 0.00369 | 0.1864 | **-1862** | **0.00239** | 0.1723 |
| S3 | -604 | 0.0081 | 0.2520 | -584 | 0.00425 | 0.0421 | -584 | 0.00474 | 0.0504 | -618 | 0.00307 | 0.0322 |
| S4 | -722 | 0.0135 | 0.4602 | -707 | 0.00710 | 0.0641 | -708 | 0.00830 | 0.0777 | -757 | 0.00595 | 0.0449 |
| S5 | -714 | 0.0293 | 0.8779 | -711 | 0.01419 | 0.1635 | -797 | 0.00644 | 0.0661 | -824 | 0.00439 | 0.1516 |
| S6 | -433 | 0.0137 | 0.3883 | -529.4 | 0.00744 | 0.0487 | -529.0 | 0.00691 | 0.0434 | -573 | 0.00488 | 0.0470 |
| S7 | -388 | 0.0308 | 0.9522 | -384 | 0.01320 | 0.1152 | -382 | 0.01233 | 0.1053 | -415 | 0.00907 | 0.0799 |
| S8 | -1073 | 0.0189 | 1.1360 | -1012.8 | 0.01439 | 0.2308 | -1013.3 | 0.01341 | 0.2128 | -1127 | 0.00940 | 0.1605 |
| S9 | -1324 | 0.0154 | 0.7718 | -1325 | 0.00540 | 0.1197 | -1340 | 0.00662 | 0.1504 | -1433 | 0.00499 | 0.0910 |
| S10 | -961 | 0.0019 | 0.0713 | -932 | 0.00096 | **0.0027** | -963 | 0.00109 | 0.0077 | -1000 | 0.00089 | 0.0076 |
| S11 | -424 | 0.0267 | 0.9003 | -445 | 0.00696 | 0.0563 | -444 | 0.00596 | 0.0511 | -467 | 0.00624 | 0.1423 |

**Table 3.**
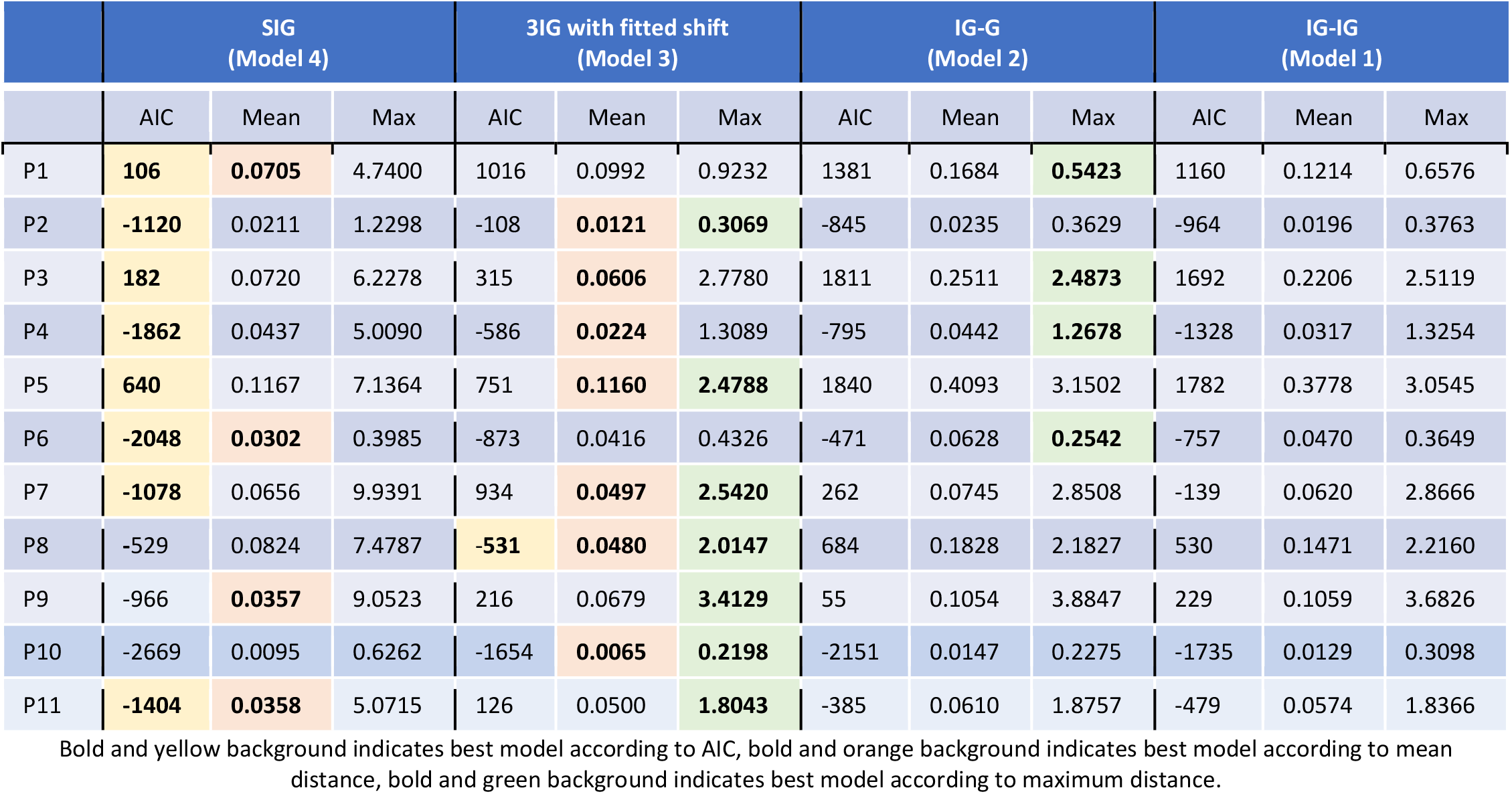
Model fit results propofol sedation cohort for Models 1-4.

|  | SIG<br>(Model 4) |  |  | 3IG with fitted shift<br>(Model 3) |  |  | IG-G<br>(Model 2) |  |  | IG-IG<br>(Model 1) |  |  |
| --- | --- | --- | --- | --- | --- | --- | --- | --- | --- | --- | --- | --- |
|  | AIC | Mean | Max | AIC | Mean | Max | AIC | Mean | Max | AIC | Mean | Max |
| P1 | **106** | **0.0705** | 4.7400 | 1016 | 0.0992 | 0.9232 | 1381 | 0.1684 | **0.5423** | 1160 | 0.1214 | 0.6576 |
| P2 | **-1120** | 0.0211 | 1.2298 | -108 | **0.0121** | **0.3069** | -845 | 0.0235 | 0.3629 | -964 | 0.0196 | 0.3763 |
| P3 | **182** | 0.0720 | 6.2278 | 315 | **0.0606** | 2.7780 | 1811 | 0.2511 | **2.4873** | 1692 | 0.2206 | 2.5119 |
| P4 | **-1862** | 0.0437 | 5.0090 | -586 | **0.0224** | 1.3089 | -795 | 0.0442 | **1.2678** | -1328 | 0.0317 | 1.3254 |
| P5 | **640** | 0.1167 | 7.1364 | 751 | **0.1160** | **2.4788** | 1840 | 0.4093 | 3.1502 | 1782 | 0.3778 | 3.0545 |
| P6 | **-2048** | **0.0302** | 0.3985 | -873 | 0.0416 | 0.4326 | -471 | 0.0628 | **0.2542** | -757 | 0.0470 | 0.3649 |
| P7 | **-1078** | 0.0656 | 9.9391 | 934 | **0.0497** | **2.5420** | 262 | 0.0745 | 2.8508 | -139 | 0.0620 | 2.8666 |
| P8 | -529 | 0.0824 | 7.4787 | **-531** | **0.0480** | **2.0147** | 684 | 0.1828 | 2.1827 | 530 | 0.1471 | 2.2160 |
| P9 | -966 | **0.0357** | 9.0523 | 216 | 0.0679 | **3.4129** | 55 | 0.1054 | 3.8847 | 229 | 0.1059 | 3.6826 |
| P10 | -2669 | 0.0095 | 0.6262 | -1654 | **0.0065** | **0.2198** | -2151 | 0.0147 | 0.2275 | -1735 | 0.0129 | 0.3098 |
| P11 | **-1404** | **0.0358** | 5.0715 | 126 | 0.0500 | **1.8043** | -385 | 0.0610 | 1.8757 | -479 | 0.0574 | 1.8366 |

**Table 4.** Model fit results propofol sedation cohort for Models 5-8. Bold and yellow background indicates best model according to AIC, bold and orange background indicates best model according to mean distance, bold and green background indicates best model according to maximum distance.

|  | 3IG with known shift<br>(Model 8) |  |  | Exponential<br>(Model 7) |  |  | Gamma<br>(Model 6) |  |  | Lognormal<br>(Model 5) |  |  |
| --- | --- | --- | --- | --- | --- | --- | --- | --- | --- | --- | --- | --- |
|  | AIC | Mean | Max | AIC | Mean | Max | AIC | Mean | Max | AIC | Mean | Max |
| P1 | 1452 | 0.4444 | 65.0399 | 562 | 0.15458 | 1.5962 | 454 | 0.09996 | 0.9653 | 222 | 0.0946 | 6.6980 |
| P2 | -718 | 0.0471 | 3.6942 | -917 | 0.02982 | 1.5101 | -915 | 0.02969 | 1.5051 | -1106 | 0.0185 | 1.3319 |
| P3 | 970 | 0.3081 | 40.4074 | 881 | 0.26527 | 14.9232 | 678 | 0.20785 | 13.3930 | 282 | 0.1187 | 7.3166 |
| P4 | 351 | 0.1384 | 29.9697 | -1056 | 0.08458 | 6.6325 | -1147 | 0.07442 | 6.3296 | -1767 | 0.0527 | 5.4607 |
| P5 | 815 | 0.3357 | 26.0754 | 1092 | 0.34848 | 18.0306 | 971 | 0.27206 | 16.1510 | 707 | 0.1471 | 8.1513 |
| P6 | -1165 | 0.1020 | 16.3274 | -1353 | 0.06431 | 1.0177 | -1443 | 0.05423 | 0.8276 | -1923 | 0.0420 | 0.6758 |
| P7 | 53 | 0.1860 | 29.9329 | 49 | 0.15824 | 13.9430 | -166 | 0.13130 | 13.1840 | -925 | 0.0875 | 10.7545 |
| P8 | -106 | 0.1273 | 12.3888 | -89 | 0.14318 | 9.5849 | -145 | 0.12754 | 9.1268 | -491 | 0.0938 | 8.5924 |
| P9 | -904 | 0.0658 | 4.4293 | -644 | 0.04718 | 9.2322 | -660 | 0.04778 | 9.3685 | **-990** | 0.0406 | 9.2512 |
| P10 | -1820 | 0.0220 | 1.8786 | -2288 | 0.01339 | 0.5838 | -2398 | 0.01205 | 0.6731 | **-2695** | 0.0091 | 0.6482 |
| P11 | -636 | 0.0970 | 7.9576 | -896 | 0.06640 | 6.0904 | -916 | 0.06031 | 5.9448 | -1335 | 0.0452 | 5.2760 |

Based on mean distance, in the awake and at rest cohort, SIG was the best model for 8 of the 11 subjects (S1, S3-S8, S11), lognormal for one subject (S2), and 3IG with fitted shift for two subjects (S9, S10). In the propofol sedation cohort, 3IG with fitted shift was the best model for 7 of the 11 subjects (P2-P5, P7, P8, P10) and SIG for the other 4 (P1, P6, P9, P11).

Based on max distance, in the awake and at rest cohort, one or more of the two difference models and the 3IG with fitted shift model were the best across 10 of the 11 subjects (more than one model performed equally well in most cases). The 3IG with fitted shift model was one of the best models for 9 of the 11 subjects (S1-S4, S6-S9, S11), the IG-G model for 5 of the 11 subjects (S3, S5-S7, S11), and the IG-IG model for 8 out of 11 subjects (S1, S3, S4, S6-S9, S11). The exponential was the best model for the remaining subject (S10). In the propofol sedation cohort, the 3IG with fitted shift was the best for 7 out of 11 subjects (P2, P5, P7-P11) and the IG-G for the remaining 4 (P1, P3, P4, P6).

Overall, all simplifications (Models 1-4) were reasonable models for the data and outperformed other distributions (Models 5-8). The 3IG with known location shift did very poorly, which suggests that the thresholding to select pulses was not interfering with the observed amplitude properties, since accounting for that threshold did not result in a well-fitting model.

Each of the simplifications of our hypothesis (Models 1-4) prioritized different aspects of the fit (Figs 2-5, S1 Appendix). The maximum distance from the 45-degree line seemed to occur at high quantiles, which was the right tail for most models. The fit in this region was prioritized by the difference models, Models 1 and 2 (IG-IG and IG-G). Models 1 and 2 fitted the right tail of the distribution by sacrificing some of the fit near the mode of the distribution, reflected in mean distance and AIC, especially as compared to SIG (Model 4). The 3IG with fitted shift (Model 3) seemed to balance the fit of both mode and tail of distribution reasonably. These results suggest that SIG was the best model in terms of efficiency, but if overall quality of fit was prioritized, the 3IG model with fitted shift may have been a better choice. The difference models (IG-IG and IG-G) were only a good choice if fit of the tail was most important.

**Fig 2.**
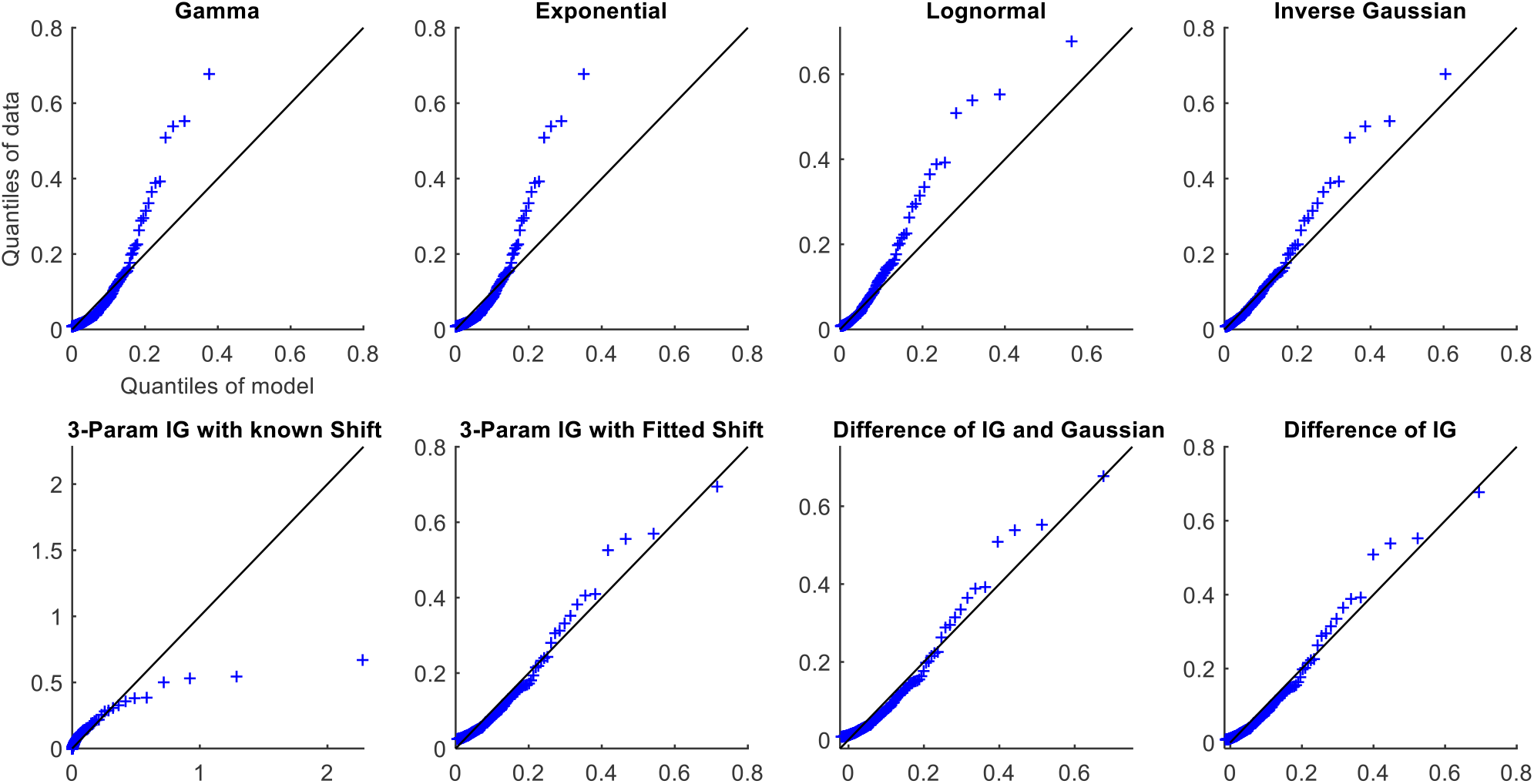
QQ plots for Subject S8 from the awake and at rest cohort.

**Fig 3.**
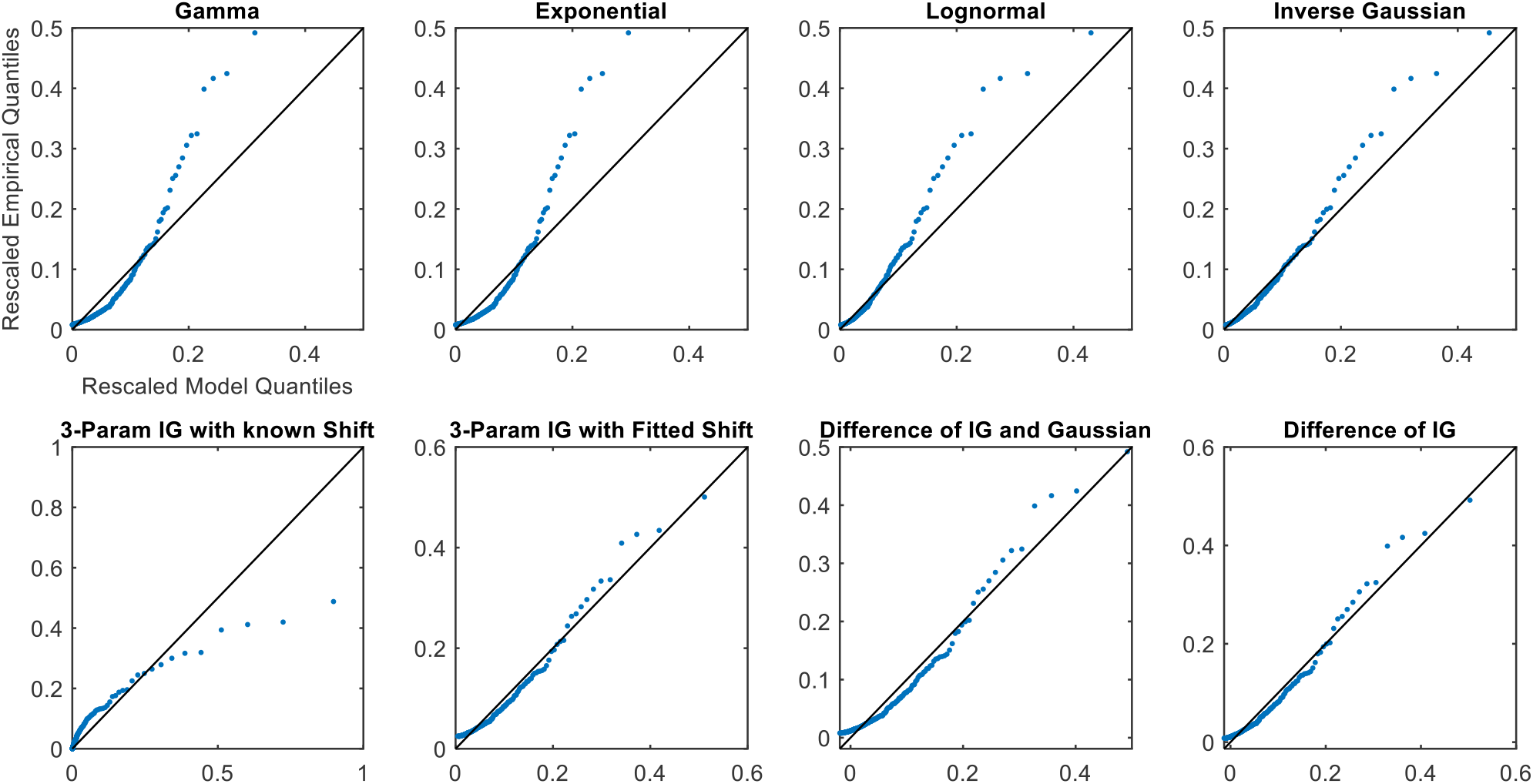
Rescaled QQ plots for Subject S8 from the awake and at rest cohort.

**Fig 4.**
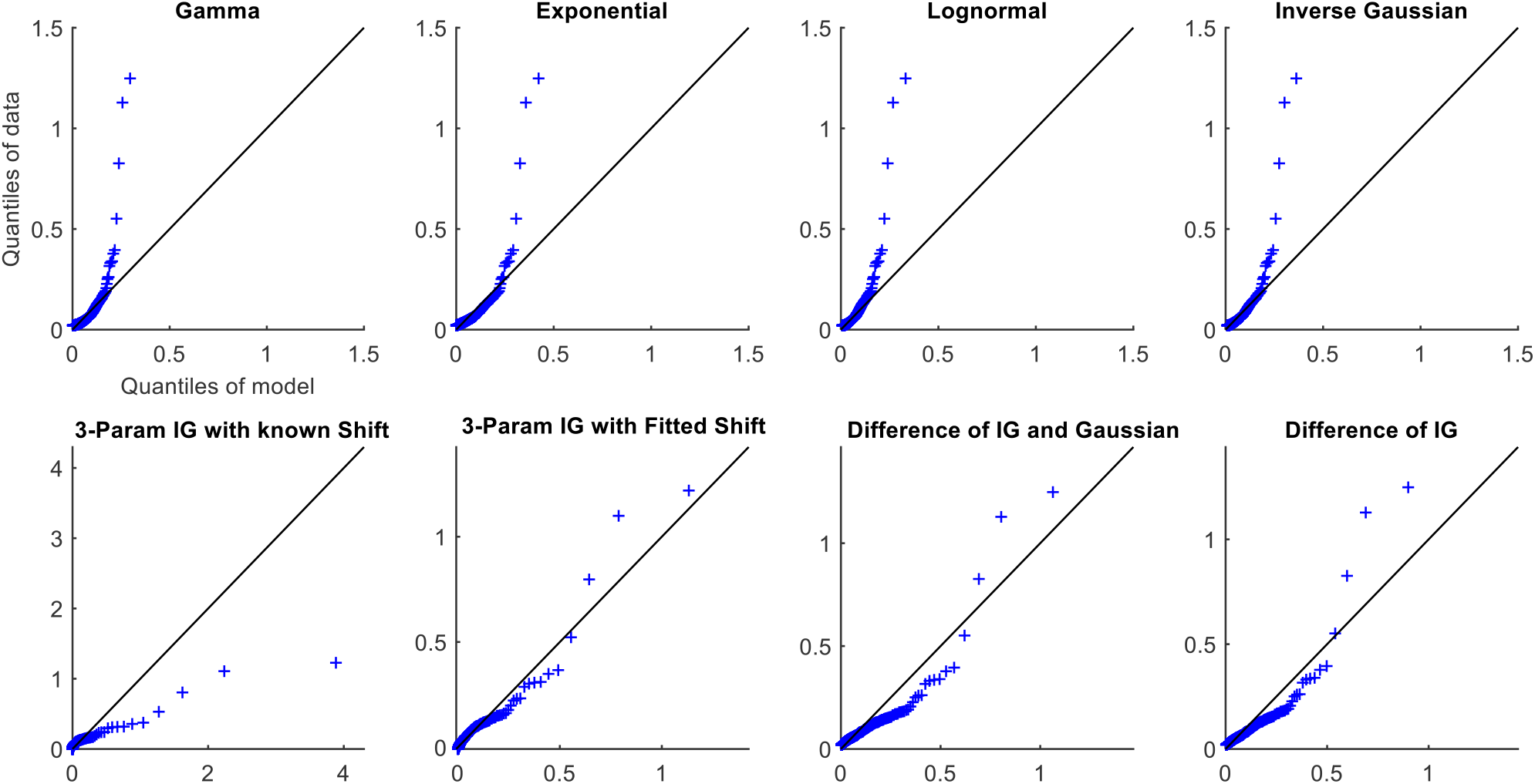
QQ plots for Subject P10 from the propofol sedation cohort.

**Fig 5.**
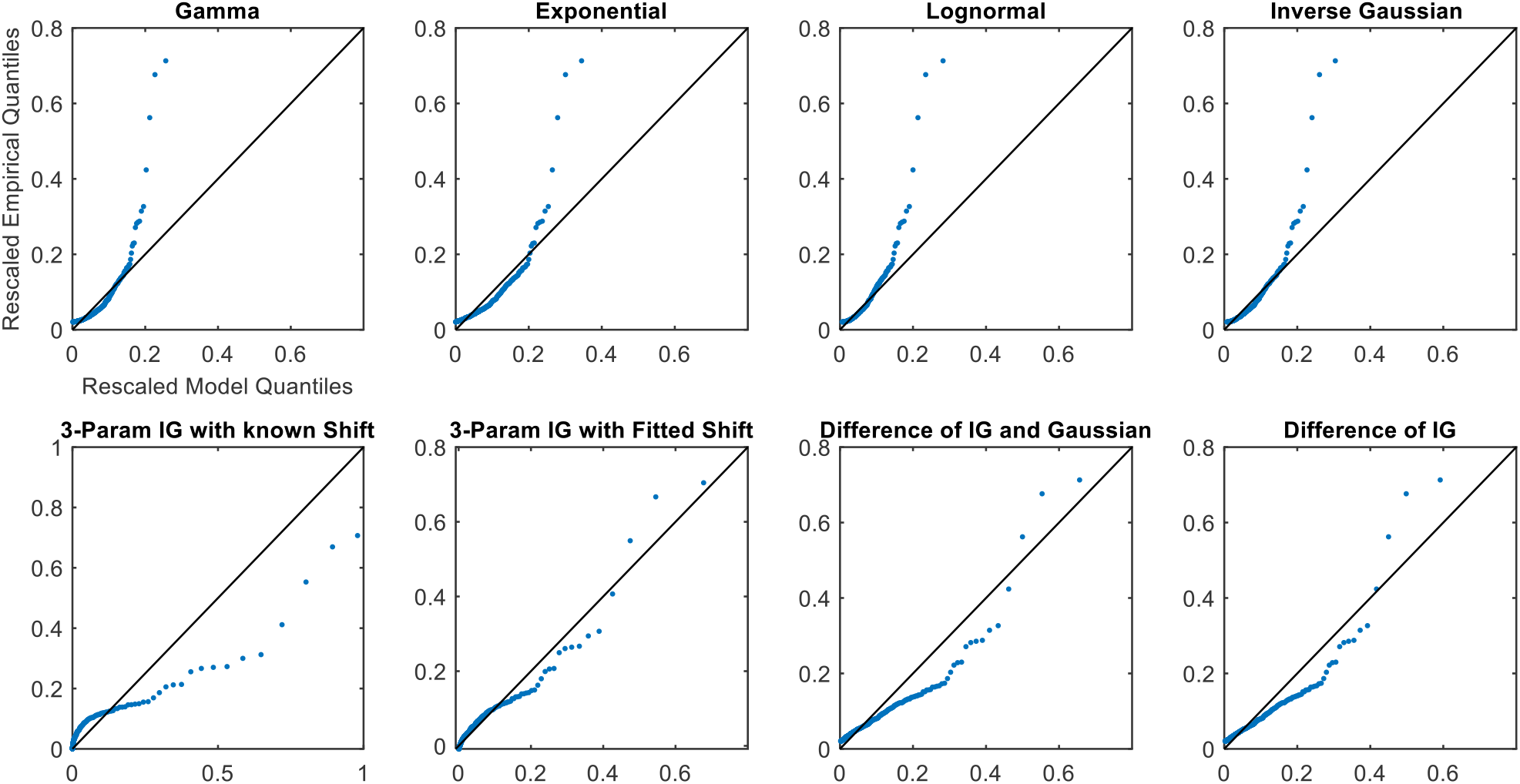
Rescaled QQ plots for Subject P10 from the propofol sedation cohort.

Finally, the parameter values (Tables 5 and 6) indicate that the progressive simplifications are logical, from the difference of two inverse Gaussians to the difference between an inverse Gaussian and a Gaussian, and then to a 3-parameter inverse Gaussian with a location shift. Across all subjects, in the IG-IG model (Model 1), the parameters of the second inverse Gaussian indicate that the ratio of 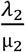 is very high, which approaches a Gaussian distribution (19). Similarly, in the IG-G model (Model 2), across all subjects, the standard deviation of the Gaussian is very small, approaching a simple point shift in the mean.

**Table 5.** Fitted parameter values for Models 1-4 for awake and at rest cohort.

|  | SIG<br>(Model 4) |  | 3IG with fitted shift<br>(Model 3) |  |  | IG-G<br>(Model 2) |  |  |  | IG-IG<br>(Model 1) |  |  |  |
| --- | --- | --- | --- | --- | --- | --- | --- | --- | --- | --- | --- | --- | --- |
| | $\mu$ | $\lambda$ | $\mu$ | $\lambda$ | $\theta$ | $\mu_1$ | $\lambda_1$ | $\mu_2$ | $\sigma_2$ | $\mu_1$ | $\lambda_1$ | $\mu_2$ | $\lambda_2$ |
| P1 | 0.5408 | 0.1373 | 0.9492 | 1.1447 | -0.4084 | 1.5151 | 3.8754 | 0.9743 | <1e-12 | 1.1246 | 1.8507 | 0.5838 | 2.6e13 |
| P2 | 0.1109 | 0.0752 | 0.0907 | 0.0145 | 0.0201 | 0.1195 | 0.0300 | 0.0086 | <1e-12 | 0.1103 | 0.0247 | 0.0000 | 1.1e-9 |
| P3 | 0.6549 | 0.1418 | 0.6616 | 0.0920 | -0.0067 | 1.0610 | 0.3000 | 0.4060 | <1e-12 | 0.9966 | 0.2592 | 0.3416 | 1.5e13 |
| P4 | 0.2179 | 0.0861 | 0.1708 | 0.0140 | 0.0471 | 0.2403 | 0.0330 | 0.0224 | <1e-12 | 0.2139 | 0.0251 | 0.0000 | 7.6e-14 |
| P5 | 0.9633 | 0.2244 | 0.9797 | 0.1560 | -0.0165 | 1.6801 | 0.6014 | 0.7169 | <1e-12 | 1.6004 | 0.5368 | 0.6371 | 2.9e13 |
| P6 | 0.1639 | 0.0615 | 0.2491 | 0.1426 | -0.0852 | 0.3366 | 0.3238 | 0.1727 | <1e-12 | 0.2694 | 0.1793 | 0.1055 | 4.8e12 |
| P7 | 0.3749 | 0.1139 | 0.2966 | 0.0216 | 0.0783 | 0.4130 | 0.0494 | 0.0381 | <1e-12 | 0.3921 | 0.0438 | 0.0172 | 7.8e11 |
| P8 | 0.3355 | 0.1294 | 0.3124 | 0.0283 | 0.0231 | 0.5568 | 0.1206 | 0.2212 | <1e-12 | 0.4894 | 0.0891 | 0.1539 | 6.9e12 |
| P9 | 0.2097 | 0.2219 | 0.0910 | 0.0019 | 0.1187 | 0.2066 | 0.0149 | <1e-10 | <1e-12 | 0.1802 | 0.0108 | 0.0000 | 8.5e-11 |
| P10 | 0.0592 | 0.1026 | 0.0302 | 0.0036 | 0.0290 | 0.0590 | 0.0203 | <1e-10 | <1e-12 | 0.0592 | 0.0262 | 0.0000 | 1.4e-14 |
| P11 | 0.2065 | 0.1057 | 0.1182 | 0.0070 | 0.0883 | 0.1462 | 0.0119 | <1e-10 | <1e-12 | 0.1547 | 0.0137 | 0.0000 | 4.8e-12 |

**Table 6.**
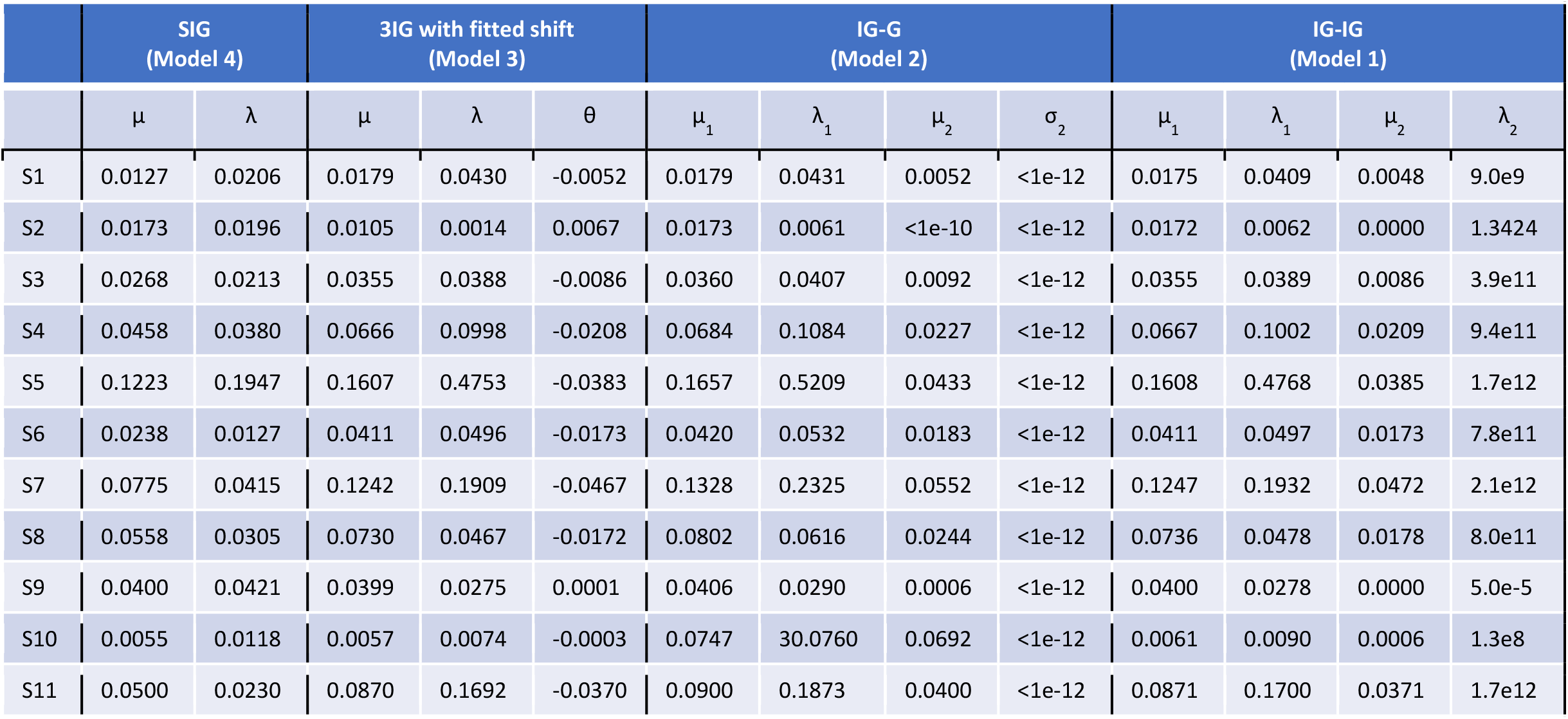
Fitted parameter values for Models 1-4 for propofol sedation cohort.

|  | SIG<br>(Model 4) |  | 3IG with fitted shift<br>(Model 3) |  |  | IG-G<br>(Model 2) |  |  |  | IG-IG<br>(Model 1) |  |  |  |
| --- | --- | --- | --- | --- | --- | --- | --- | --- | --- | --- | --- | --- | --- |
| | $\mu$ | $\lambda$ | $\mu$ | $\lambda$ | $\theta$ | $\mu_1$ | $\lambda_1$ | $\mu_2$ | $\sigma_2$ | $\mu_1$ | $\lambda_1$ | $\mu_2$ | $\lambda_2$ |
| S1 | 0.0127 | 0.0206 | 0.0179 | 0.0430 | -0.0052 | 0.0179 | 0.0431 | 0.0052 | <1e-12 | 0.0175 | 0.0409 | 0.0048 | 9.0e9 |
| S2 | 0.0173 | 0.0196 | 0.0105 | 0.0014 | 0.0067 | 0.0173 | 0.0061 | <1e-10 | <1e-12 | 0.0172 | 0.0062 | 0.0000 | 1.3424 |
| S3 | 0.0268 | 0.0213 | 0.0355 | 0.0388 | -0.0086 | 0.0360 | 0.0407 | 0.0092 | <1e-12 | 0.0355 | 0.0389 | 0.0086 | 3.9e11 |
| S4 | 0.0458 | 0.0380 | 0.0666 | 0.0998 | -0.0208 | 0.0684 | 0.1084 | 0.0227 | <1e-12 | 0.0667 | 0.1002 | 0.0209 | 9.4e11 |
| S5 | 0.1223 | 0.1947 | 0.1607 | 0.4753 | -0.0383 | 0.1657 | 0.5209 | 0.0433 | <1e-12 | 0.1608 | 0.4768 | 0.0385 | 1.7e12 |
| S6 | 0.0238 | 0.0127 | 0.0411 | 0.0496 | -0.0173 | 0.0420 | 0.0532 | 0.0183 | <1e-12 | 0.0411 | 0.0497 | 0.0173 | 7.8e11 |
| S7 | 0.0775 | 0.0415 | 0.1242 | 0.1909 | -0.0467 | 0.1328 | 0.2325 | 0.0552 | <1e-12 | 0.1247 | 0.1932 | 0.0472 | 2.1e12 |
| S8 | 0.0558 | 0.0305 | 0.0730 | 0.0467 | -0.0172 | 0.0802 | 0.0616 | 0.0244 | <1e-12 | 0.0736 | 0.0478 | 0.0178 | 8.0e11 |
| S9 | 0.0400 | 0.0421 | 0.0399 | 0.0275 | 0.0001 | 0.0406 | 0.0290 | 0.0006 | <1e-12 | 0.0400 | 0.0278 | 0.0000 | 5.0e-5 |
| S10 | 0.0055 | 0.0118 | 0.0057 | 0.0074 | -0.0003 | 0.0747 | 30.0760 | 0.0692 | <1e-12 | 0.0061 | 0.0090 | 0.0006 | 1.3e8 |
| S11 | 0.0500 | 0.0230 | 0.0870 | 0.1692 | -0.0370 | 0.0900 | 0.1873 | 0.0400 | <1e-12 | 0.0871 | 0.1700 | 0.0371 | 1.7e12 |

## DISCUSSION

EDA consists of two simultaneous components or processes at different timescales, the tonic and phasic. Within phasic EDA, there are two sources of relevant physiological characteristics, the timing (temporal information) and size of pulses (amplitudes). In our previous work, we developed a point process model for the temporal information (2,3). In this paper, we present a model for pulse amplitudes.

We used EDA data from two subject cohorts, a set of healthy volunteers while awake and at rest and another set of healthy volunteers under controlled propofol sedation, to verify our hypothesis that the pulse amplitudes in EDA were characterized by highly regular statistical structure. This statistical structure was consistent with the integrate-and-fire physiology that describes sweat gland function and accounted for the effect of the background, in addition to stimulus intensity, on amplitude. We fitted eight models to the pulse amplitudes in EDA, four of which were simplifications to varying degrees of our hypothesis that each pulse can be modeled as the difference of inverse Gaussians. We quantified the goodness-of-fit with three methods: AIC, and mean and maximum distances from the 45-degree line on the QQ plot. Together, we showed that the model fits were consistent with not only integrate-and-fire sweat gland physiology, but also the combined effects of varying stimulus intensity and EDA background on the dynamics of generated pulses.

The different simplifications of our hypothesis each emphasized fitting different parts of the distribution. For example, the two difference models, Models 1 and 2 (IG-IG and IG-G), both prioritized fitting the right tail of the distribution (the largest pulses) by sacrificing some of the fit near the mode of the distribution. In contrast, the SIG model (Model 4) prioritized fitting the mode of the distribution. The 3IG with fitted shift model balanced both. In the visualization, this can be seen as the varying slope of the QQ-plot relative to the 45-degree line. Going from the most simplified (SIG) to the least simplified (IG-IG) model (from Model 4 to Model 1), each is affected progressively more in terms of slope by the largest pulses. This occurs because the additional parameters, whether the location shift or those of the subtracted distribution, allowed the model to better tailor the fit of the tail.

There were some interesting differences in the performances of the models between the two subject cohorts. In the awake and at rest cohort, the SIG model (Model 4) performed best in terms of AIC and mean distance from the diagonal, while the other simplification models (Models 1-3) all performed well by maximum distance from the diagonal (the 3IG with fitted shift perhaps doing the best). In contrast, in the propofol sedation cohort, the SIG (Model 4) was the best only according to AIC, while the 3IG with fitted shift was the best in terms of both mean and maximum distance from the diagonal. This may reflect a difference in dynamics between both cohorts. Perhaps the more simplified model performed better in the awake and at rest cohort because there were fewer changing dynamics when subjects were largely at rest compared to a changing concentration of drug with known autonomic effects. Or alternatively, perhaps a longer duration of data in the propofol sedation cohort contained more dynamics that required additional complexity in the model.

The result of this study creates a direct link between the physiology of sweat glands and the statistical structure of the pulse amplitude data collected at the skin surface. The most detailed of existing models of EDA are founded in signal processing methods alone, are computationally complex, and make assumptions about pulse amplitudes out of necessity (6-10). However, looking to the physiology provided a principled framework by which to propose low-order models for pulse amplitudes (maximum of 4 parameters) that account for the effects of both stimulus and background. This result has implications for understanding and tracking the sympathetic component of the autonomic nervous system in a more meaningful way, including both temporal and amplitude information from EDA.

In future work, we will use this result to robustly and accurately capture the valuable physiological characteristics from both the timing and amplitude of pulses in any EDA dataset. We will study the dynamics of both the timing and amplitudes of pulses over time, applying history dependent inverse Gaussian models like those developed for heart rate variability (26-29) and methods for marked point processes (20,21). We will also study EDA in other contexts, such as during sleep, with pain, and under general anesthesia. Eventually, these methods will have both clinical and non-clinical applications, such as in emotional state and stress detection. Our findings provide a principled, physiologically based approach for extending EDA analyses to these more complex and important applications.

## ACKNOWLEDGMENTS

We would like to thank the MIT Clinical Research Center Staff. This work was partially funded by funds from the Picower Institute for Learning and Memory, the National Science Foundation Graduate Research Fellowship Program, the MIT Office of Graduate Education, and NIH Award P01-GM118629 (to E.N.B.).

## SUPPORTING INFORMATION CAPTIONS

**S1 Appendix. Additional figures for all subjects**.

**S2 Appendix. Densities for Models 3-8**.

**S3 Appendix. Derivation of method of moments estimates for Models 1 and 2**.

