## Supplementary material for "Elementary Integrate-and-Fire Process Underlies Pulse Amplitudes in Electrodermal Activity": S1 Appendix

### AWAKE AND AT REST COHORT

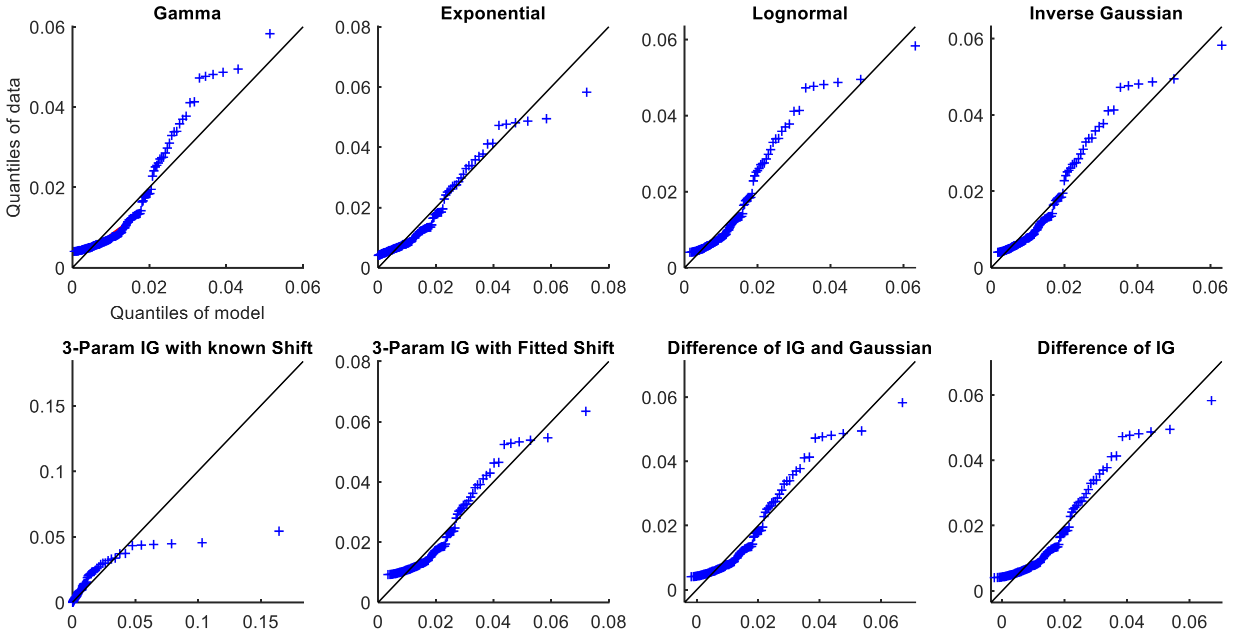


Fig. S1. QQ plots for Subject S1 from the awake and at rest cohort

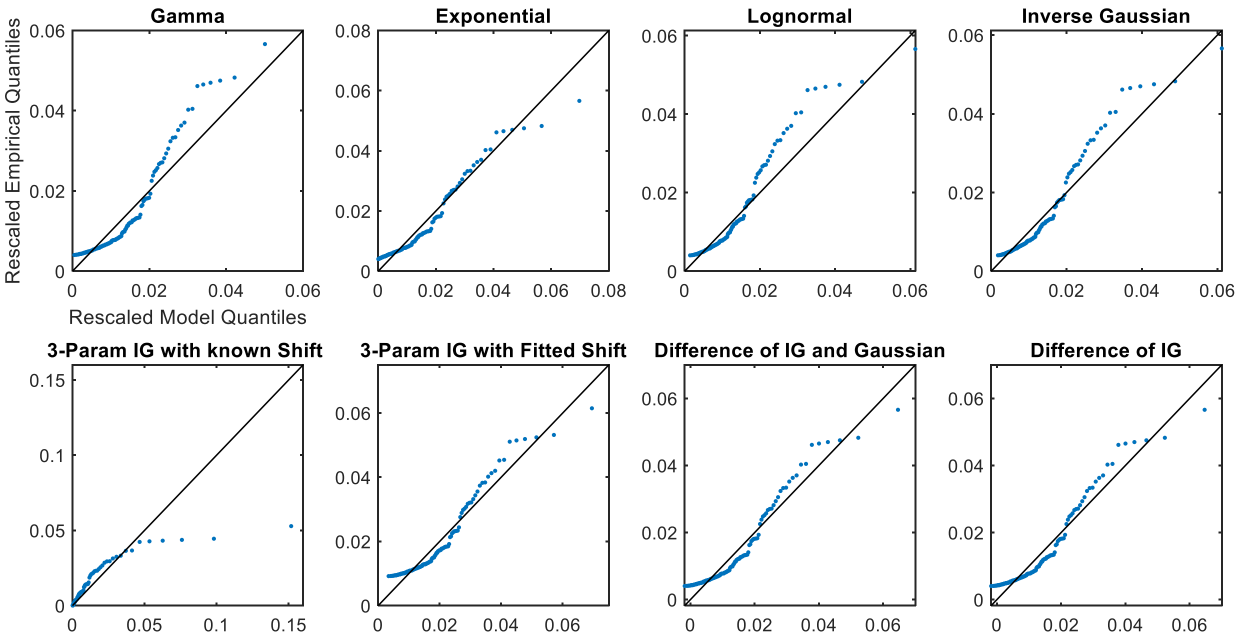


Fig. S2. Rescaled QQ plots for Subject S1 from the awake and at rest cohort

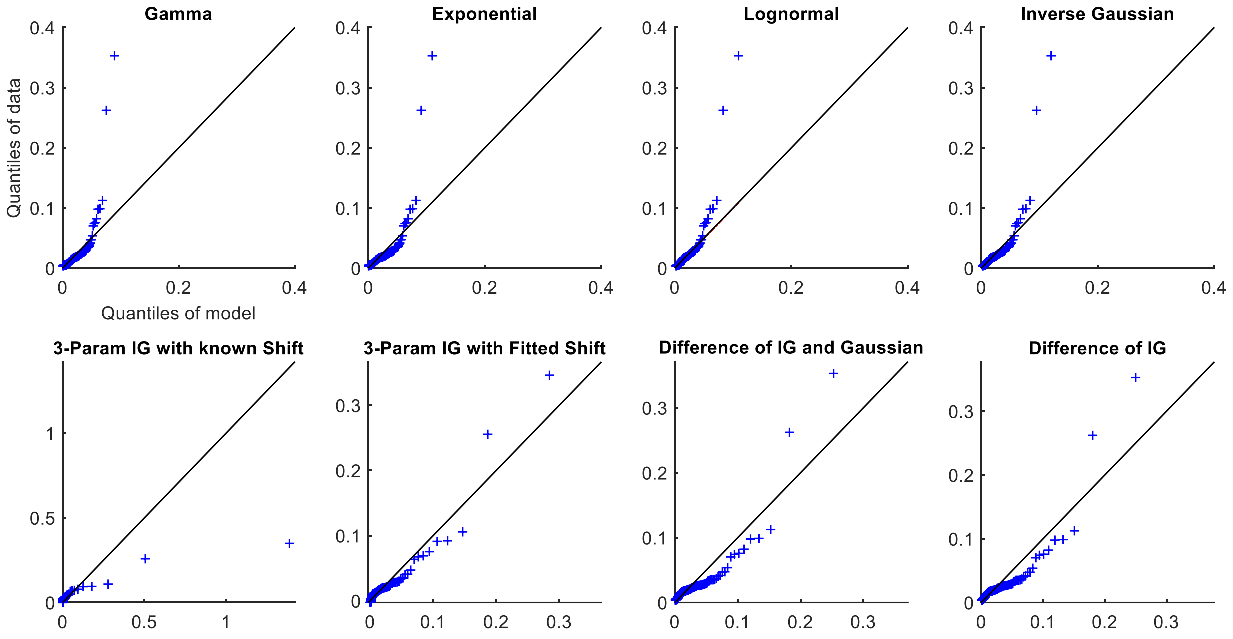


Fig. S3. QQ plots for Subject S2 from the awake and at rest cohort

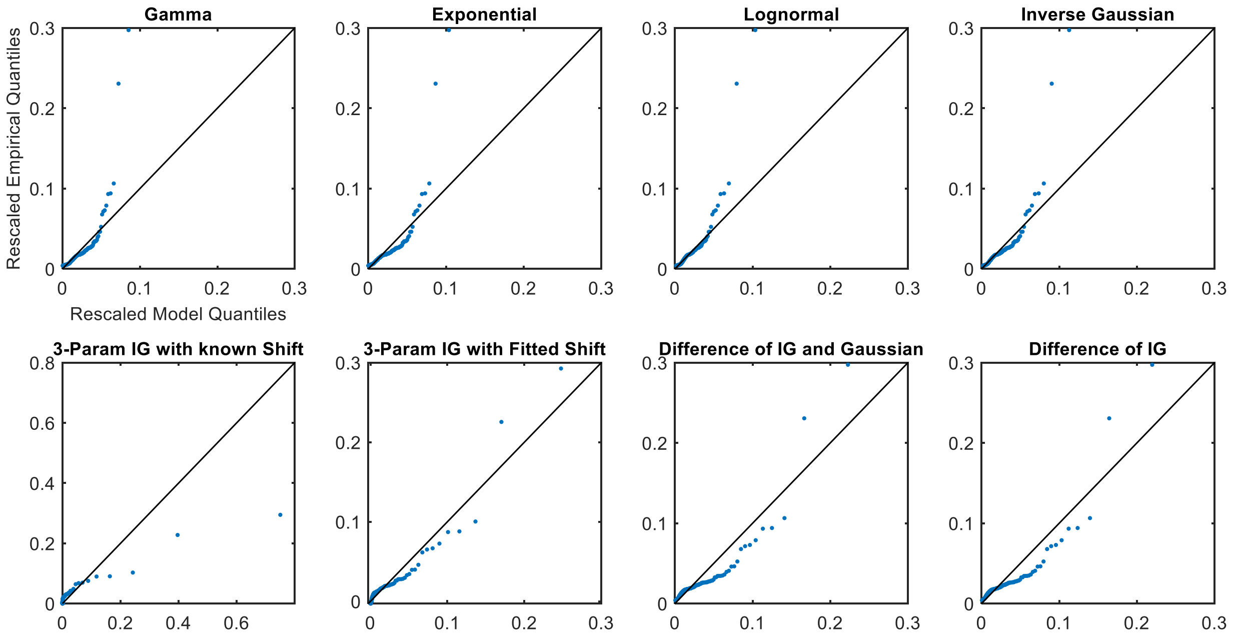


Fig. S4. Rescaled QQ plots for Subject S2 from the awake and at rest cohort

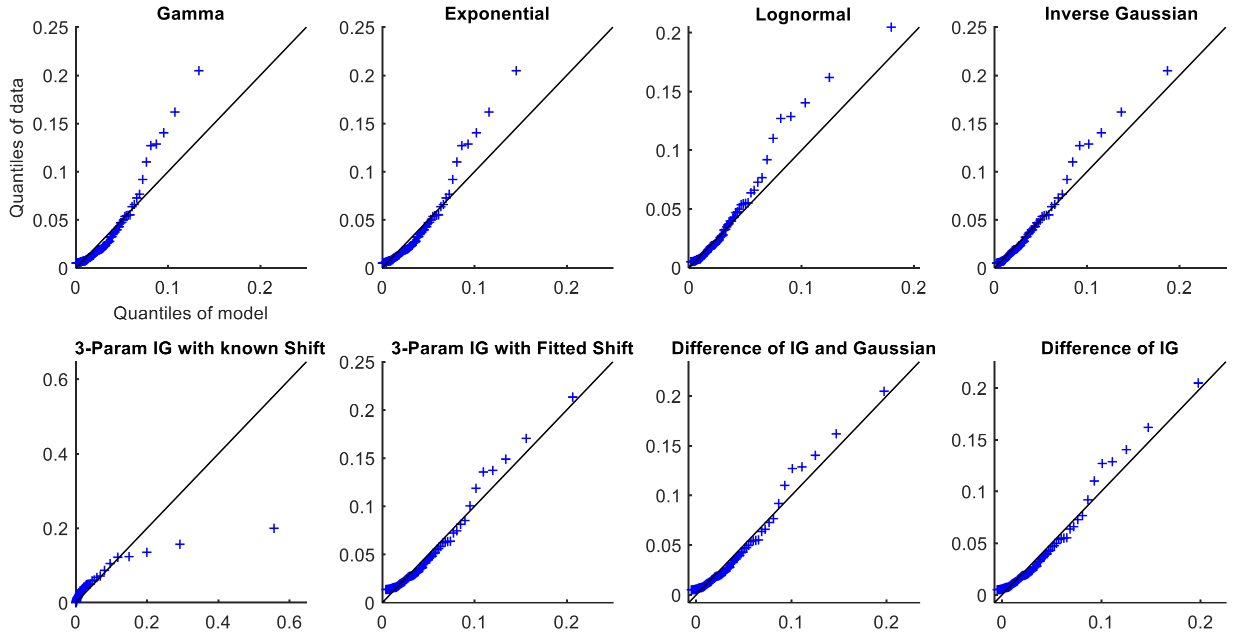


Fig. S5. QQ plots for Subject S3 from the awake and at rest cohort

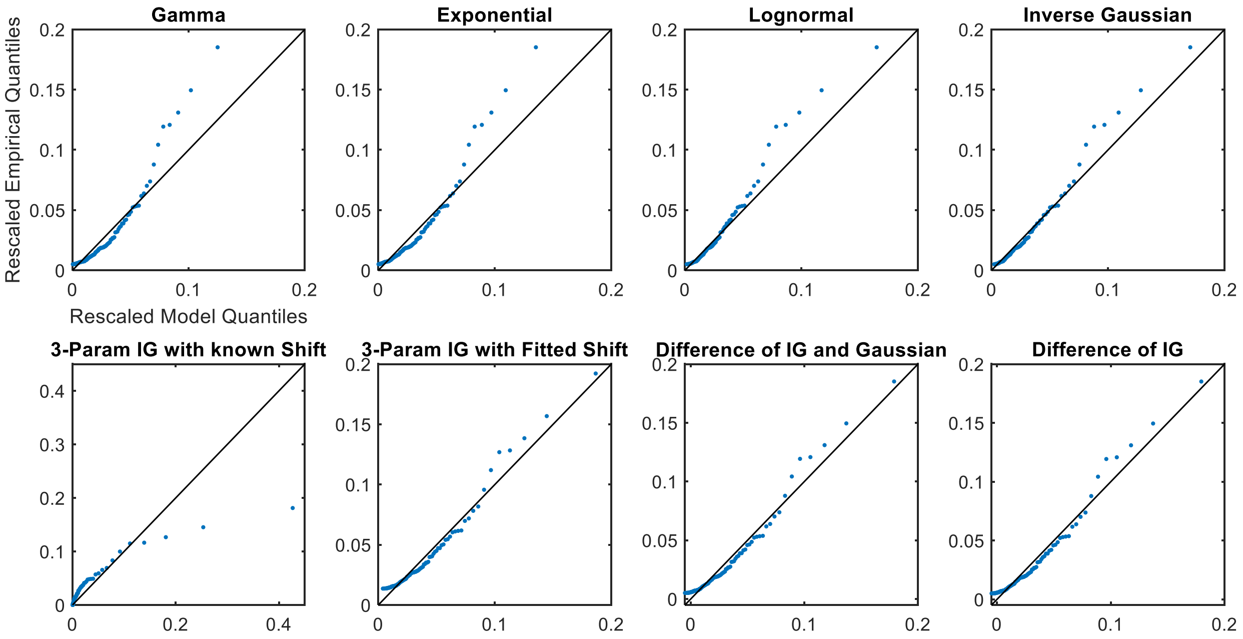


Fig. S6. Rescaled QQ plots for Subject S3 from the awake and at rest cohort

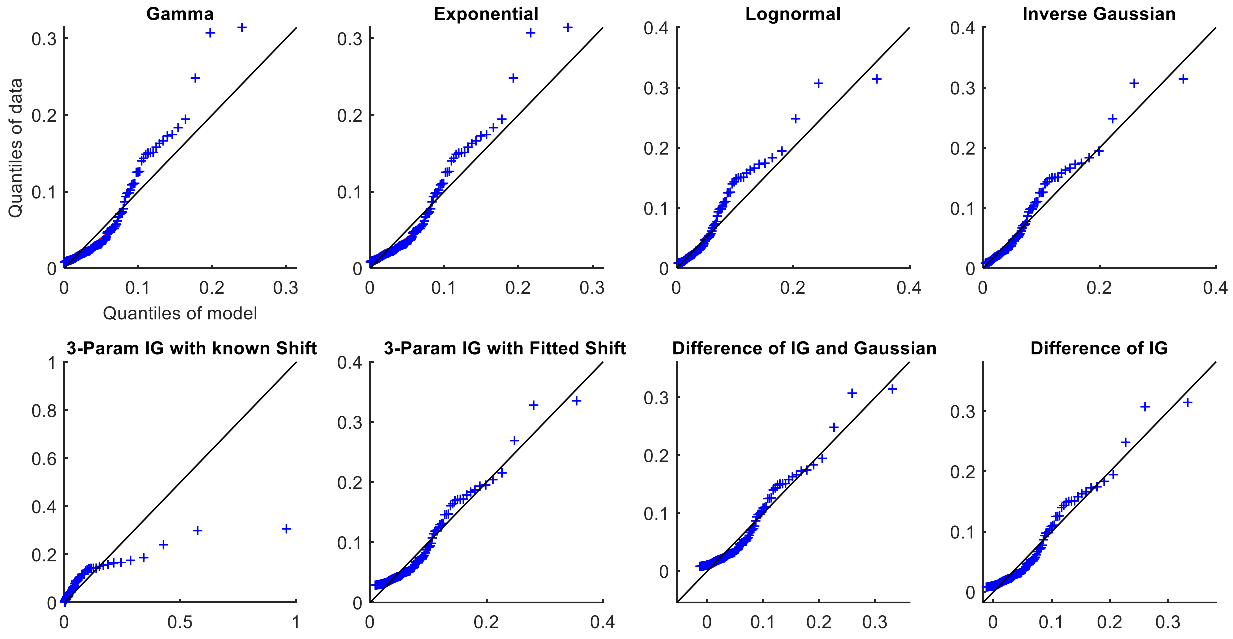


Fig. S7. QQ plots for Subject S4 from the awake and at rest cohort

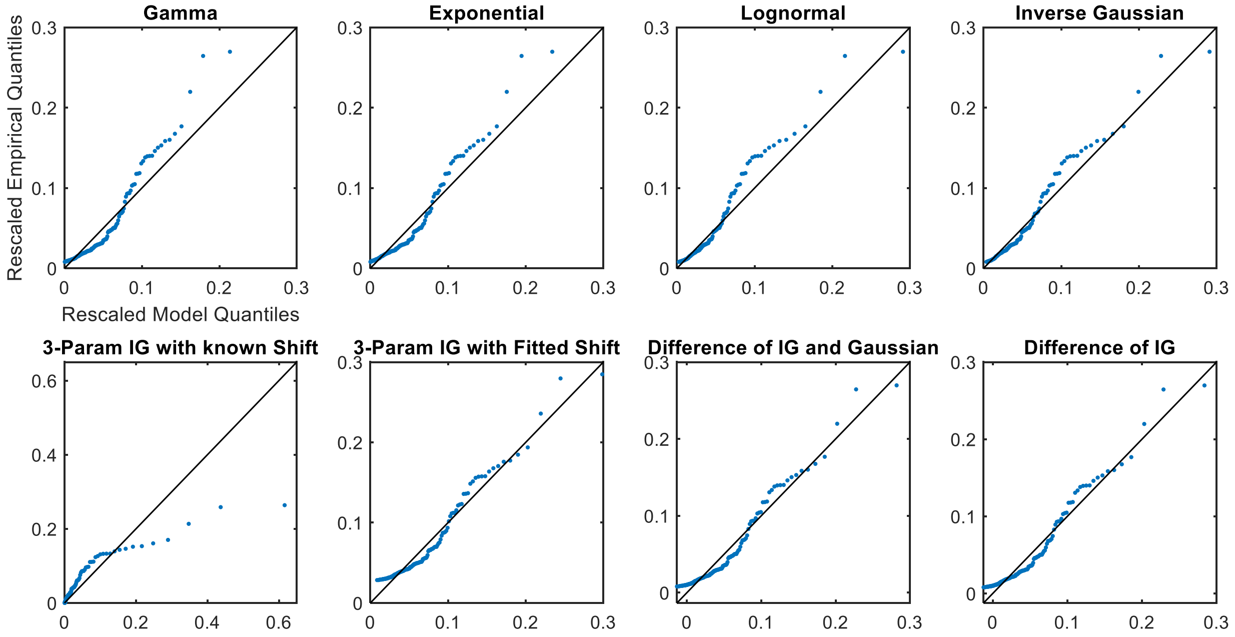


Fig. S8. Rescaled QQ plots for Subject S4 from the awake and at rest cohort

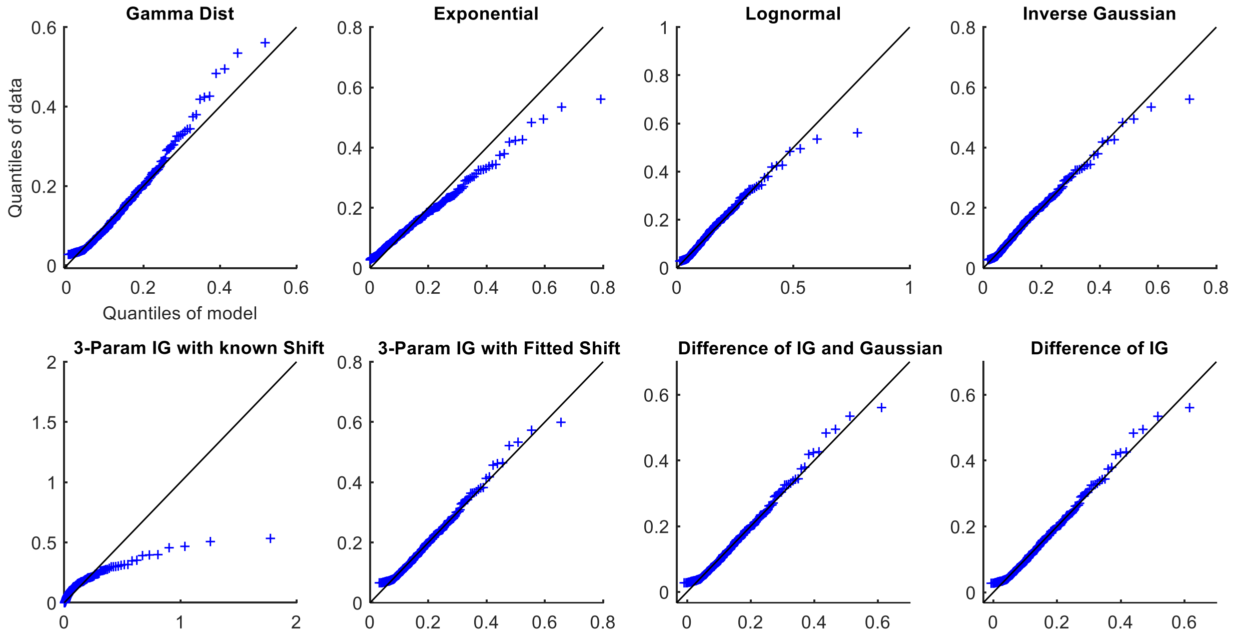


Fig. S9. QQ plots for Subject S5 from the awake and at rest cohort

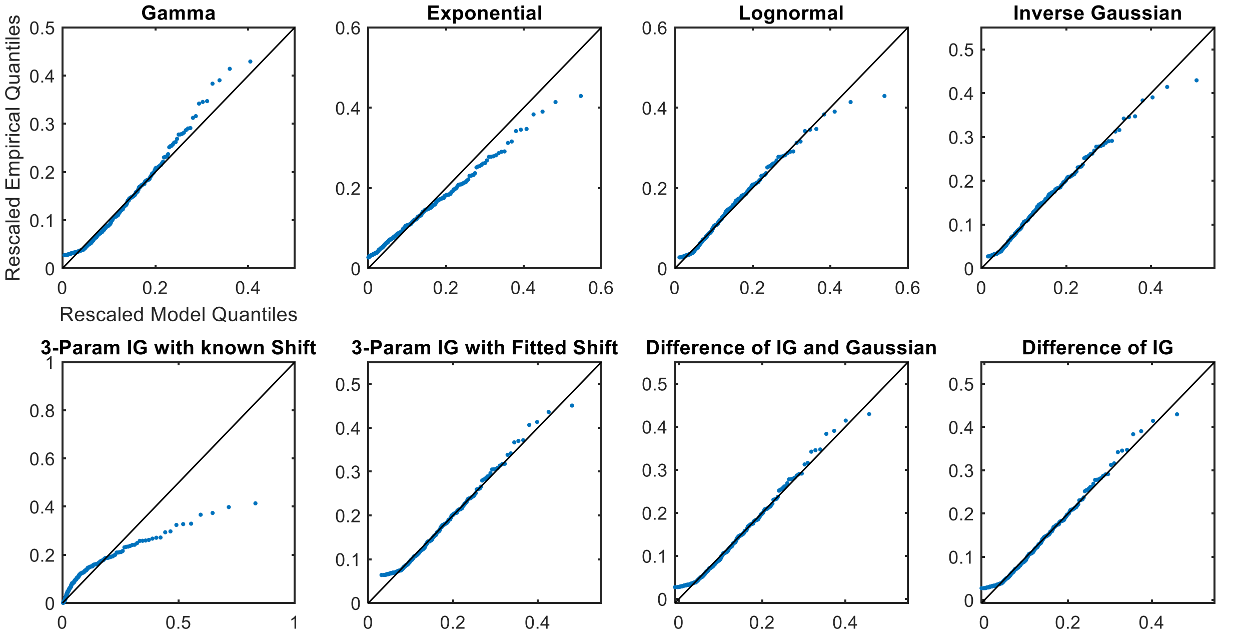


Fig. S10. Rescaled QQ plots for Subject S5 from the awake and at rest cohort

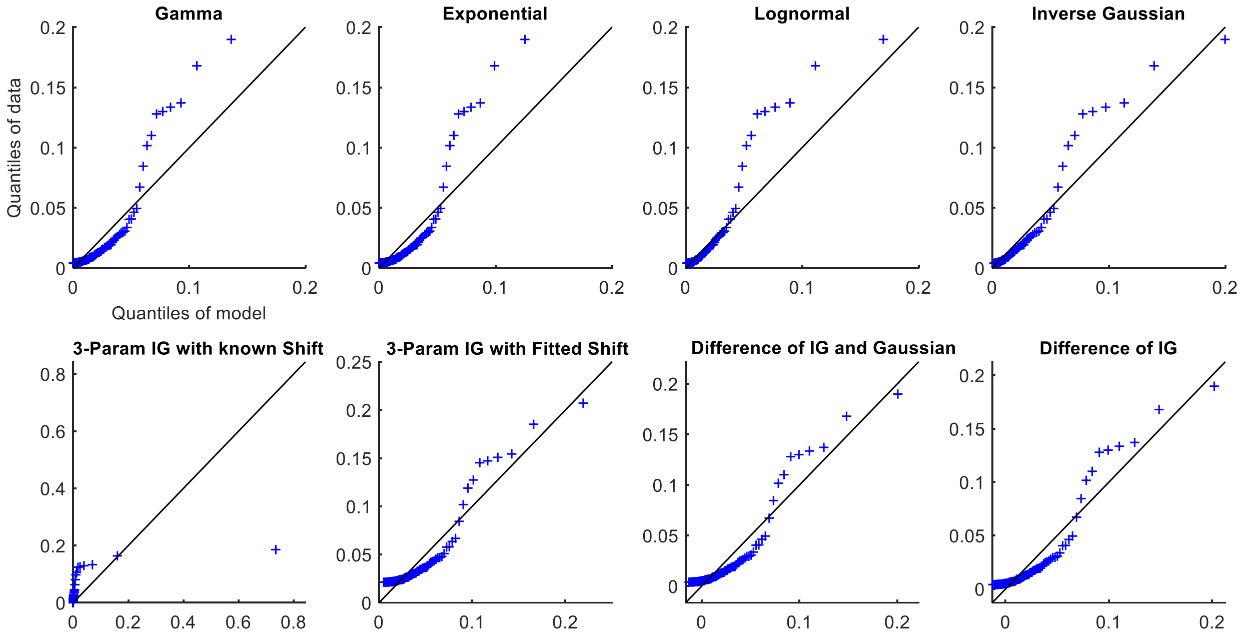


Fig. S11. QQ plots for Subject S6 from the awake and at rest cohort

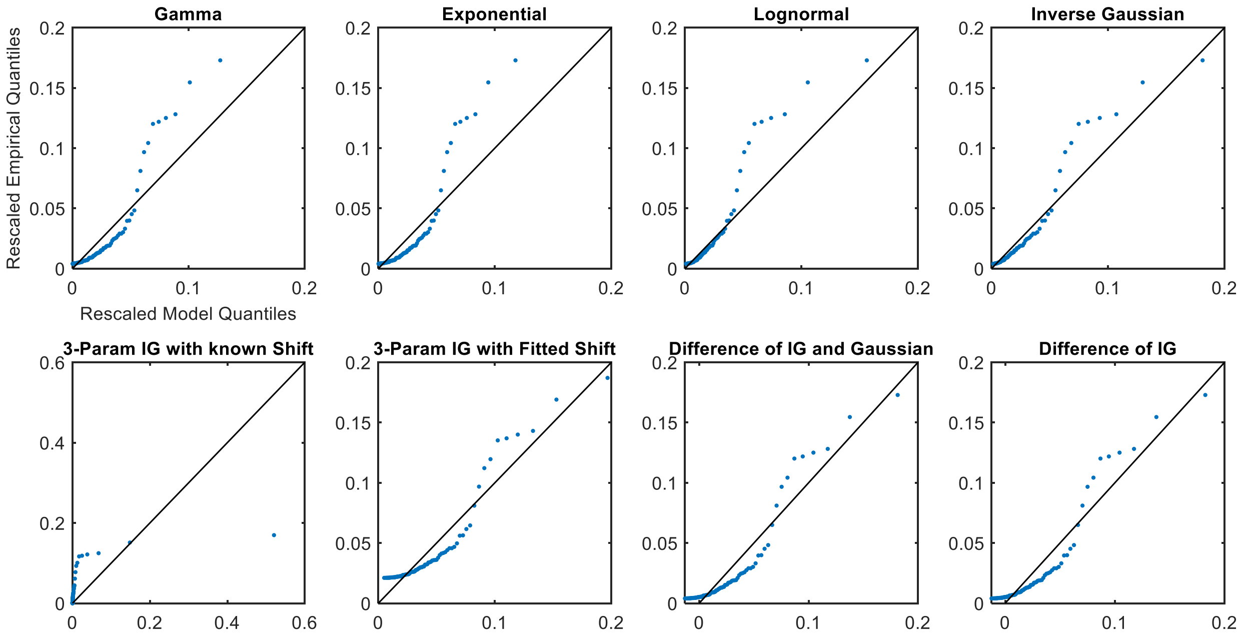


Fig. S12. Rescaled QQ plots for Subject S6 from the awake and at rest cohort

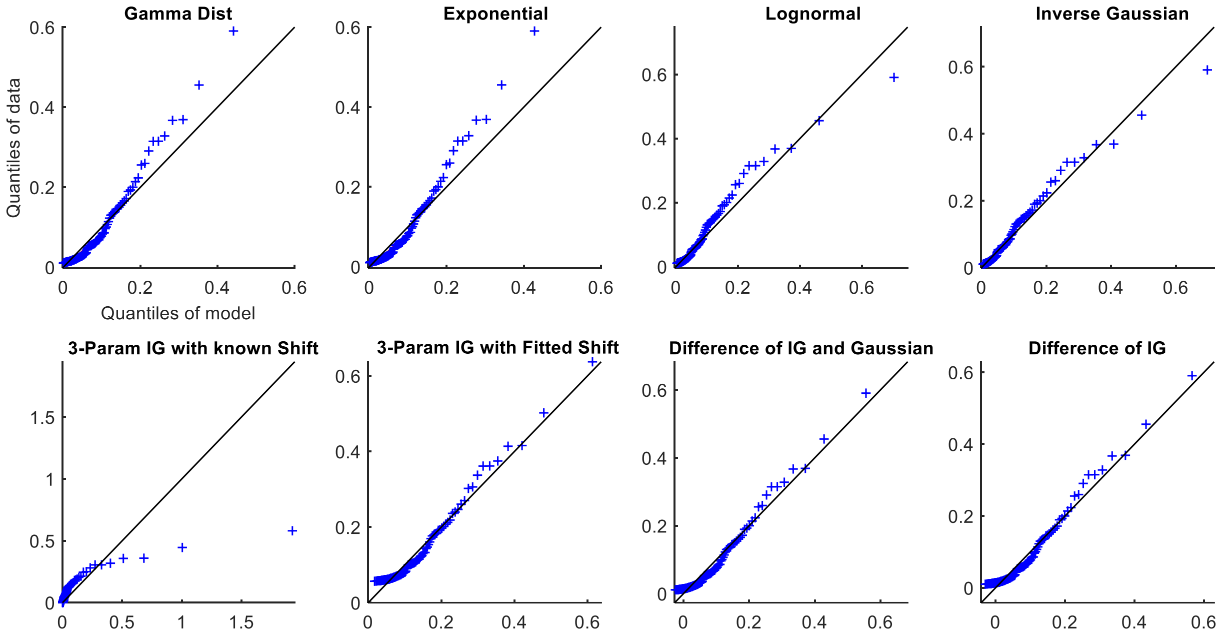


Fig. S13. QQ plots for Subject S7 from the awake and at rest cohort

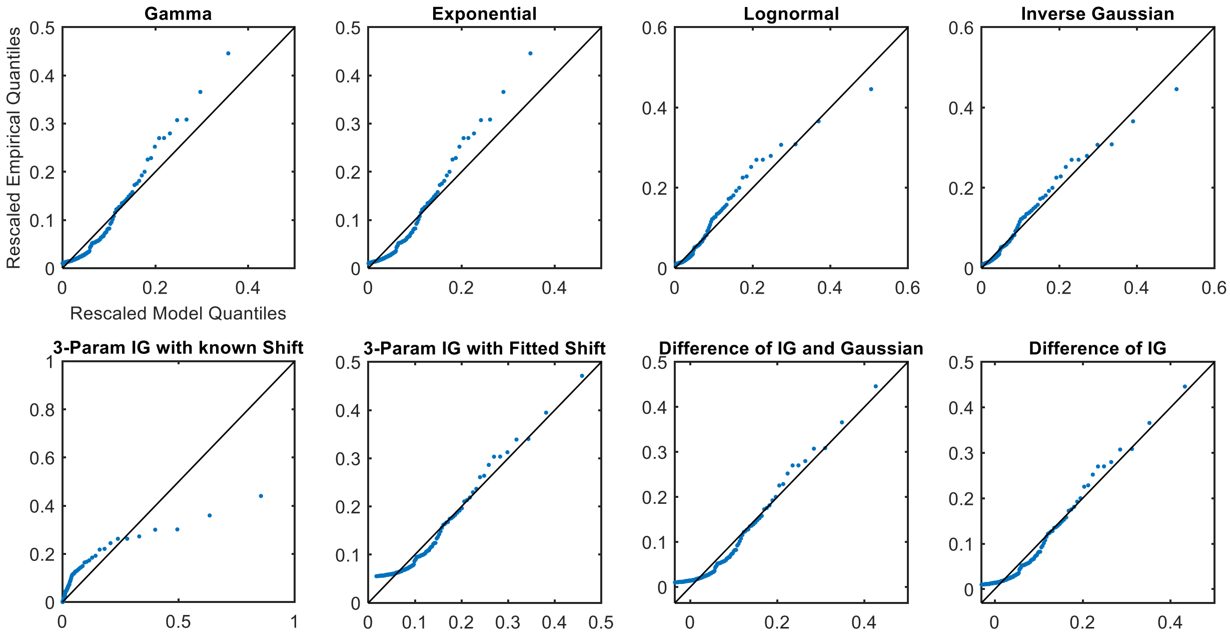


Fig. S14. Rescaled QQ plots for Subject S7 from the awake and at rest cohort

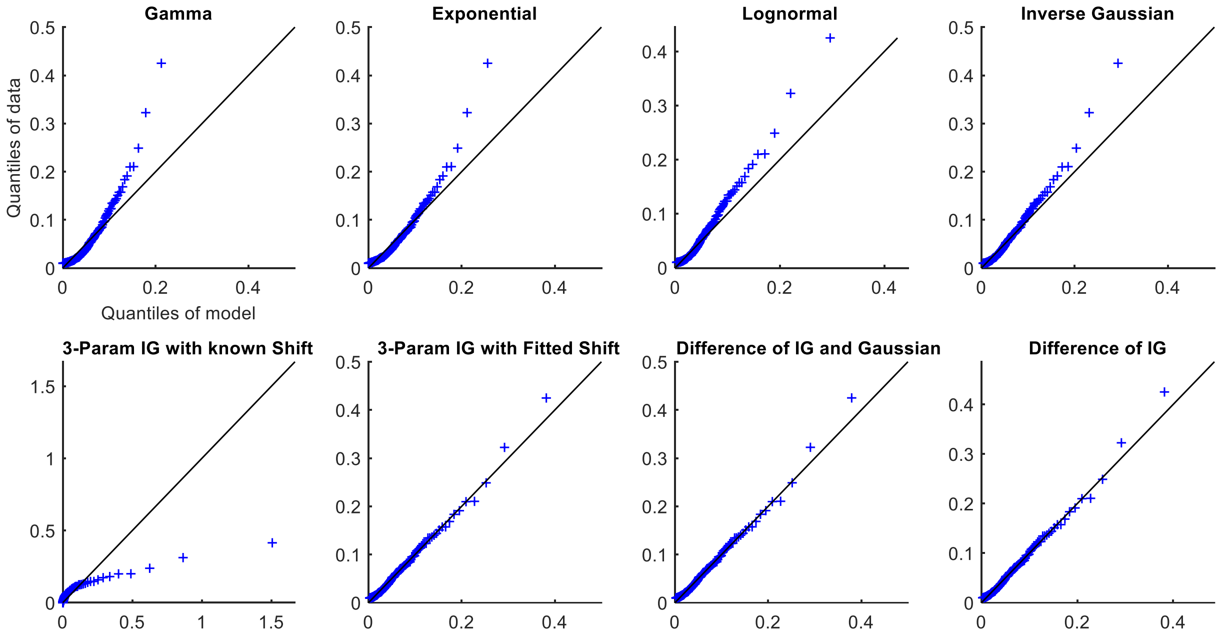


Fig. S15. QQ plots for Subject S9 from the awake and at rest cohort

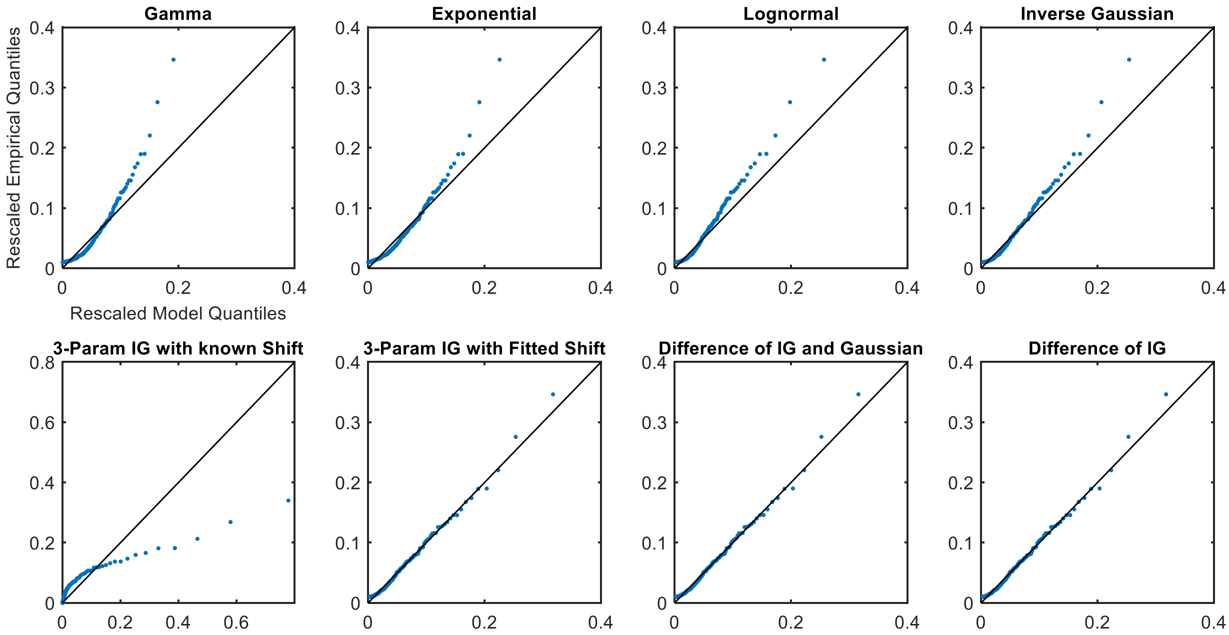


Fig. S16. Rescaled QQ plots for Subject S9 from the awake and at rest cohort

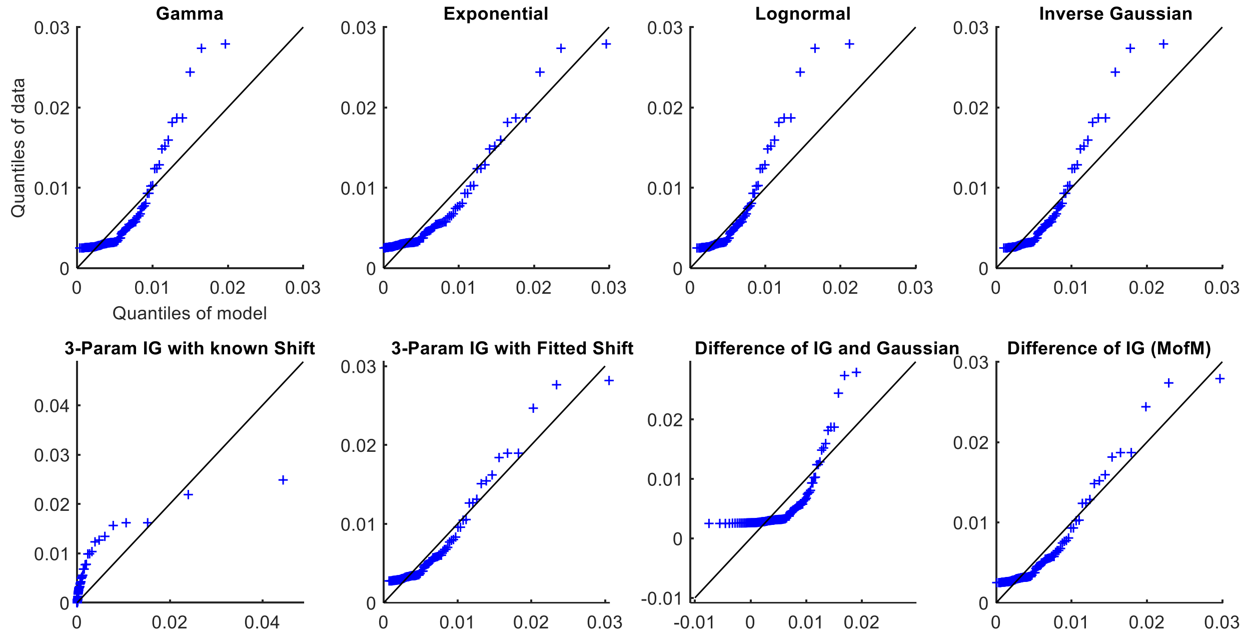


Fig. S17. QQ plots for Subject S10 from the awake and at rest cohort

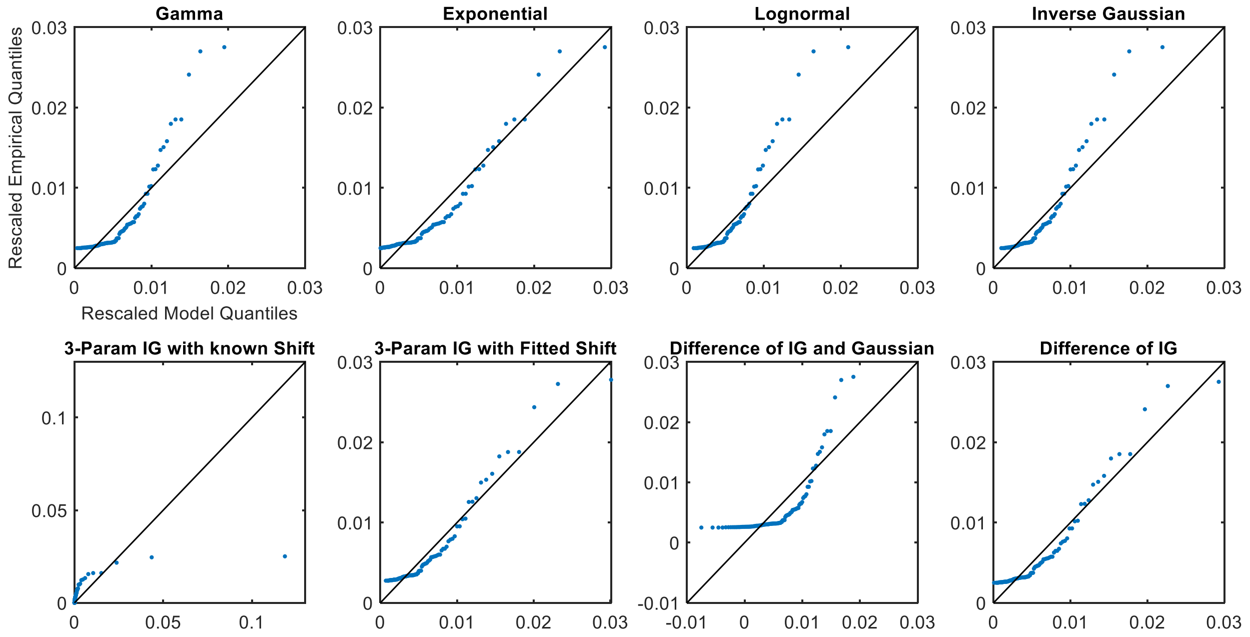


Fig. S18. Rescaled QQ plots for Subject S10 from the awake and at rest cohort

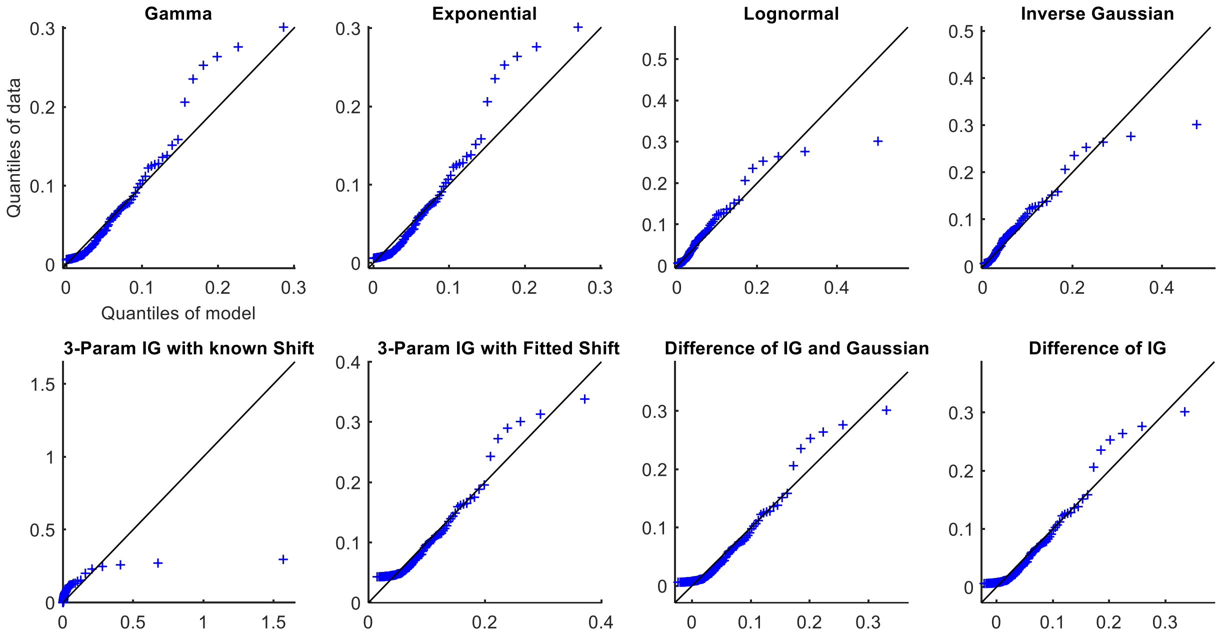


Fig. S19. QQ plots for Subject S11 from the awake and at rest cohort

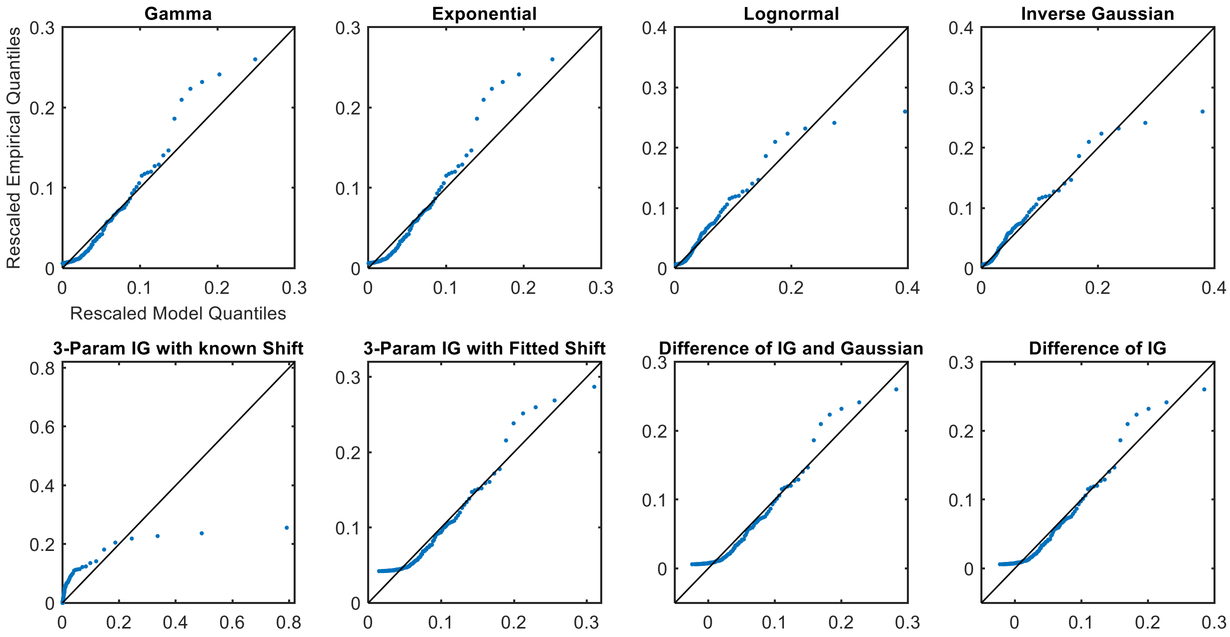


Fig. S20. Rescaled QQ plots for Subject S11 from the awake and at rest cohort

### PROPOFOL SEDATION COHORT

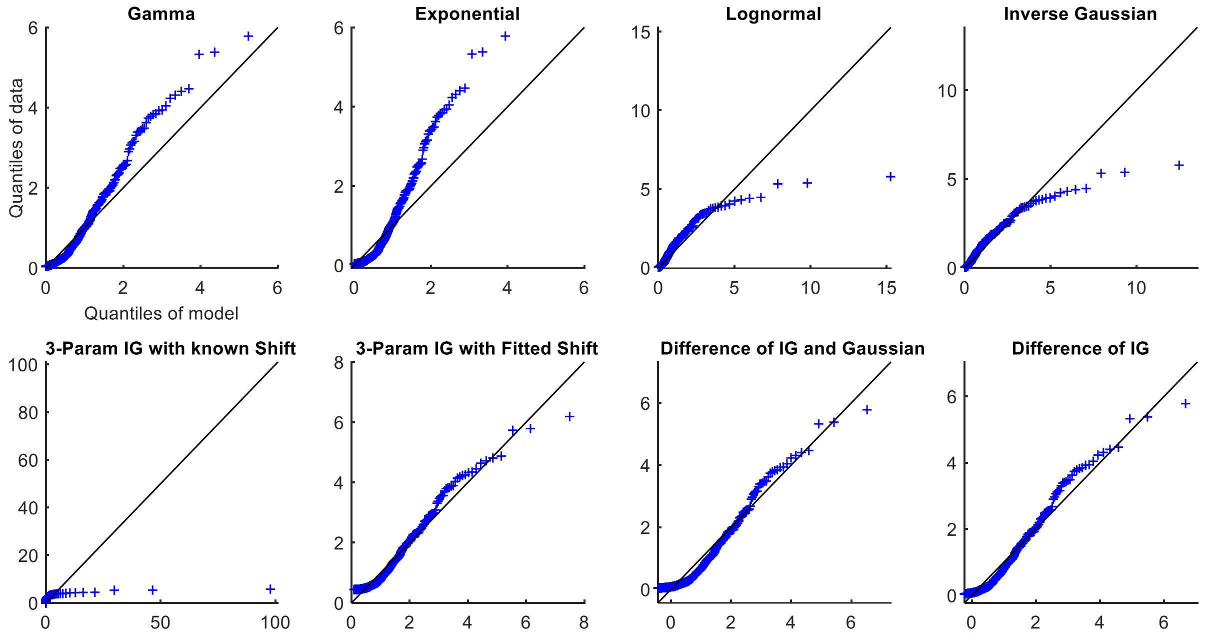


Fig. S21. QQ plots for Subject P1 from the propofol sedation cohort

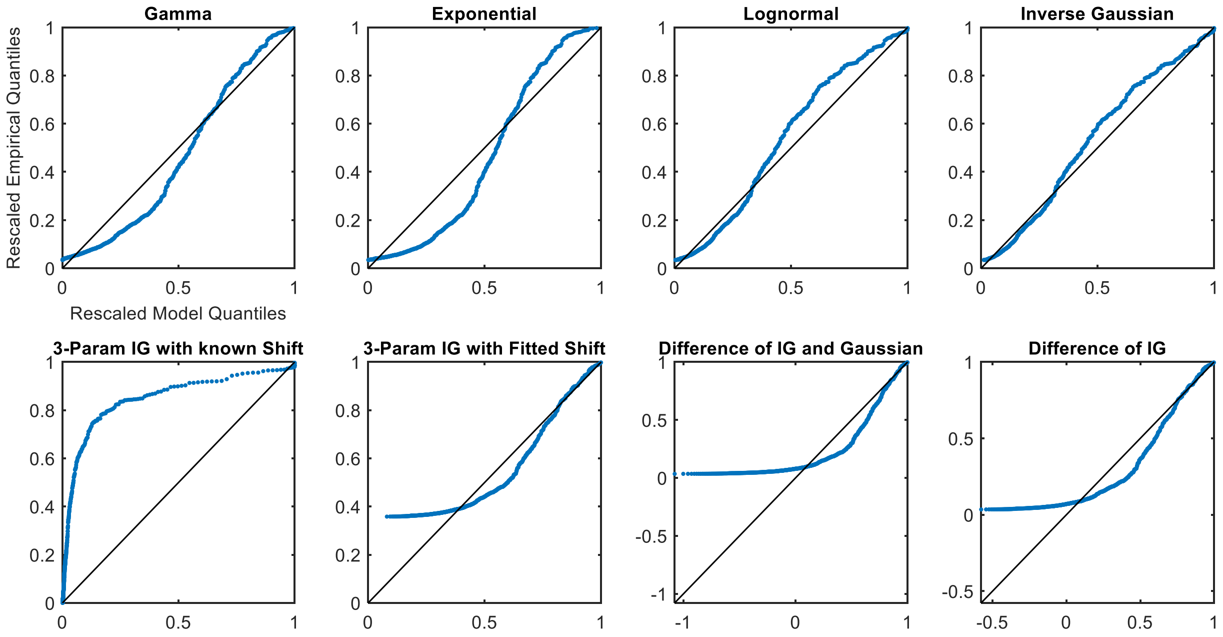


Fig. S22. Rescaled QQ plots for Subject P1 from the propofol sedation cohort

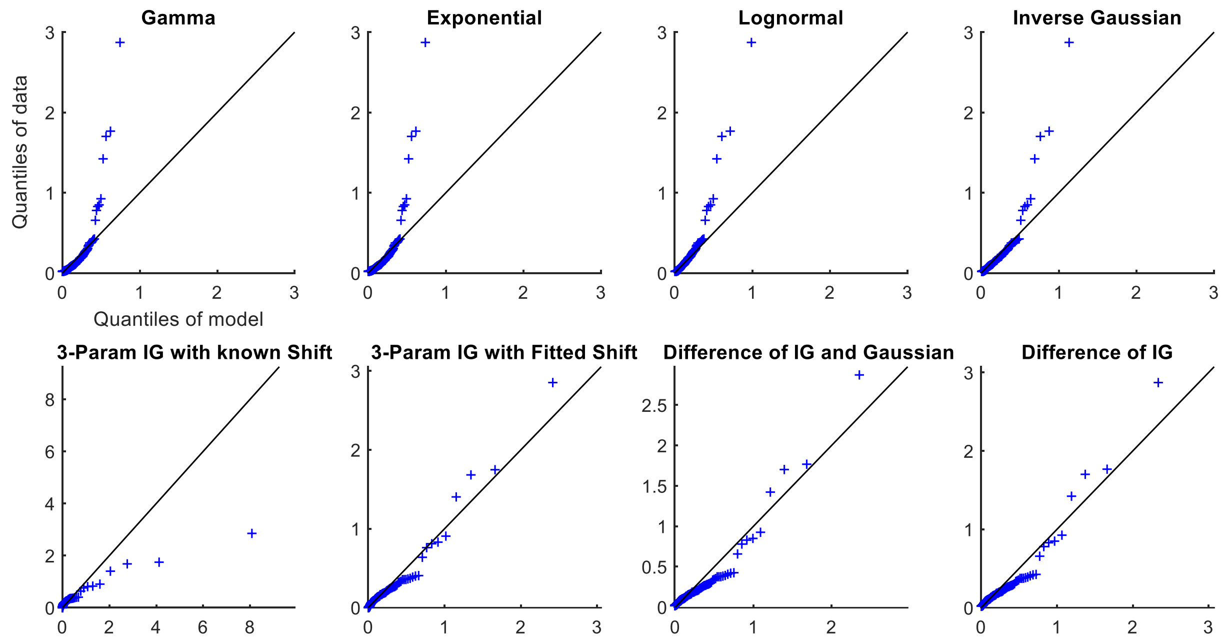


Fig. S23. QQ plots for Subject P2 from the propofol sedation cohort

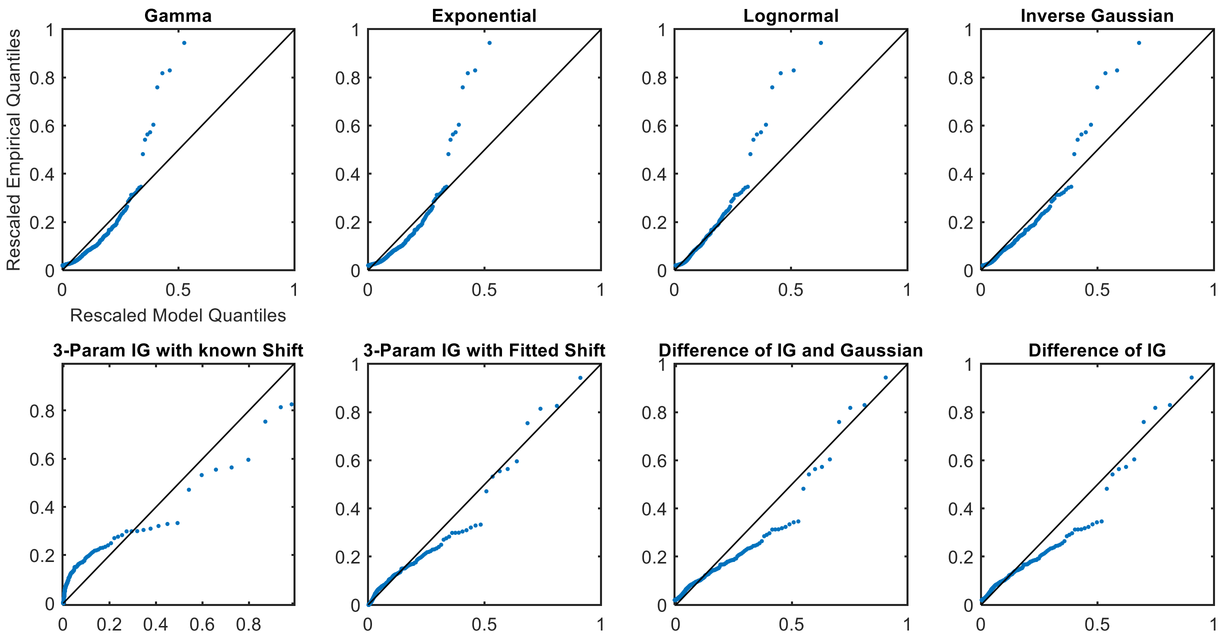


Fig. S24. Rescaled QQ plots for Subject P2 from the propofol sedation cohort

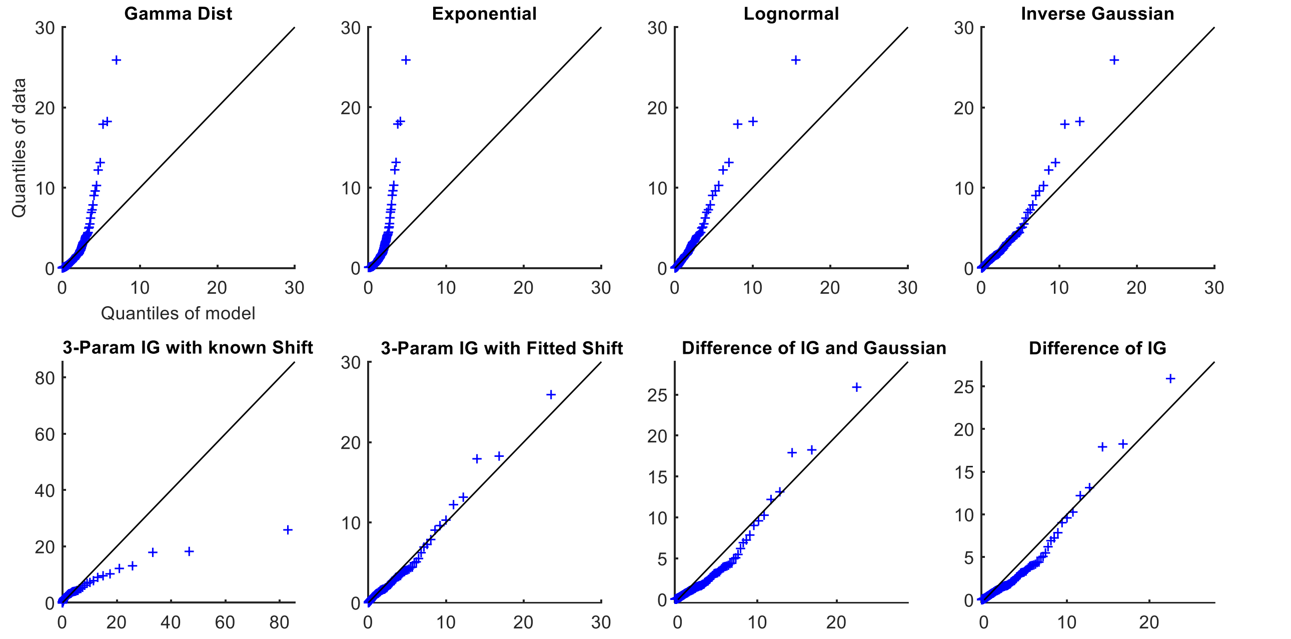


Fig. S25. QQ plots for Subject P3 from the propofol sedation cohort

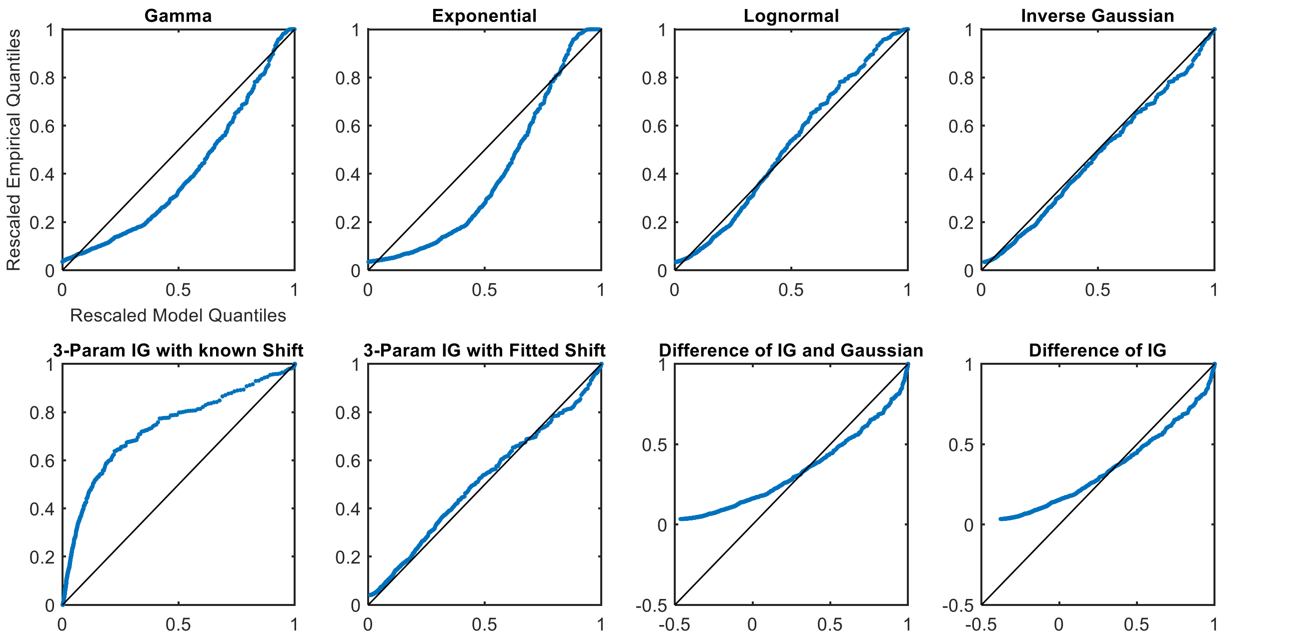


Fig. S26. Rescaled QQ plots for Subject P3 from the propofol sedation cohort

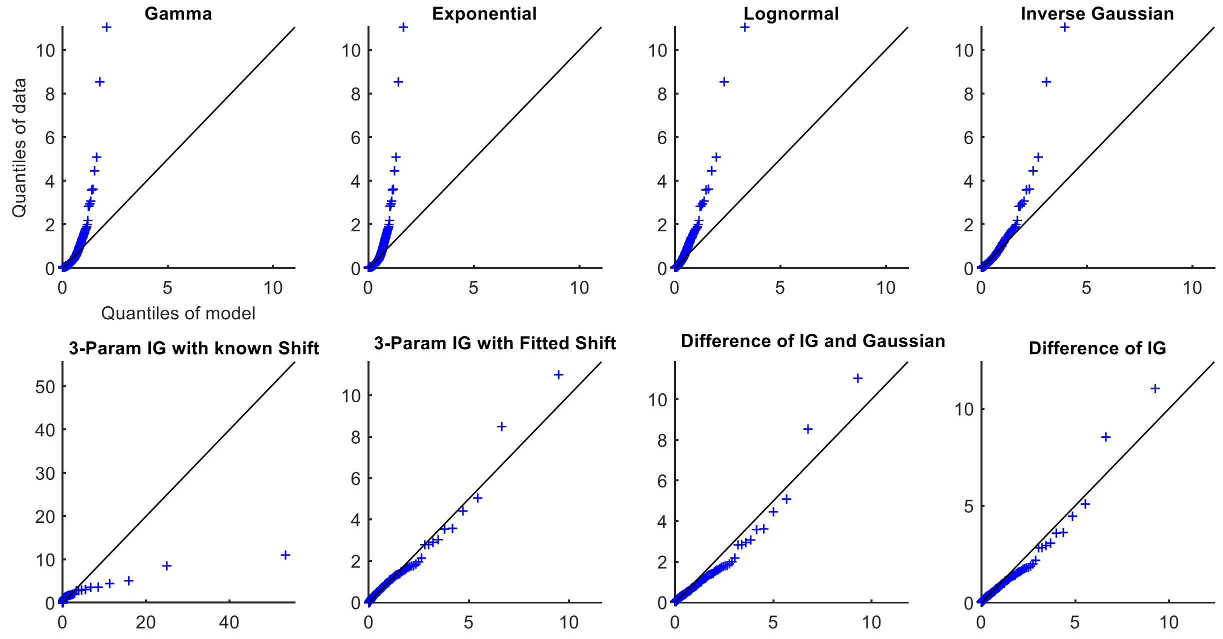


Fig. S27. QQ plots for Subject P4 from the propofol sedation cohort

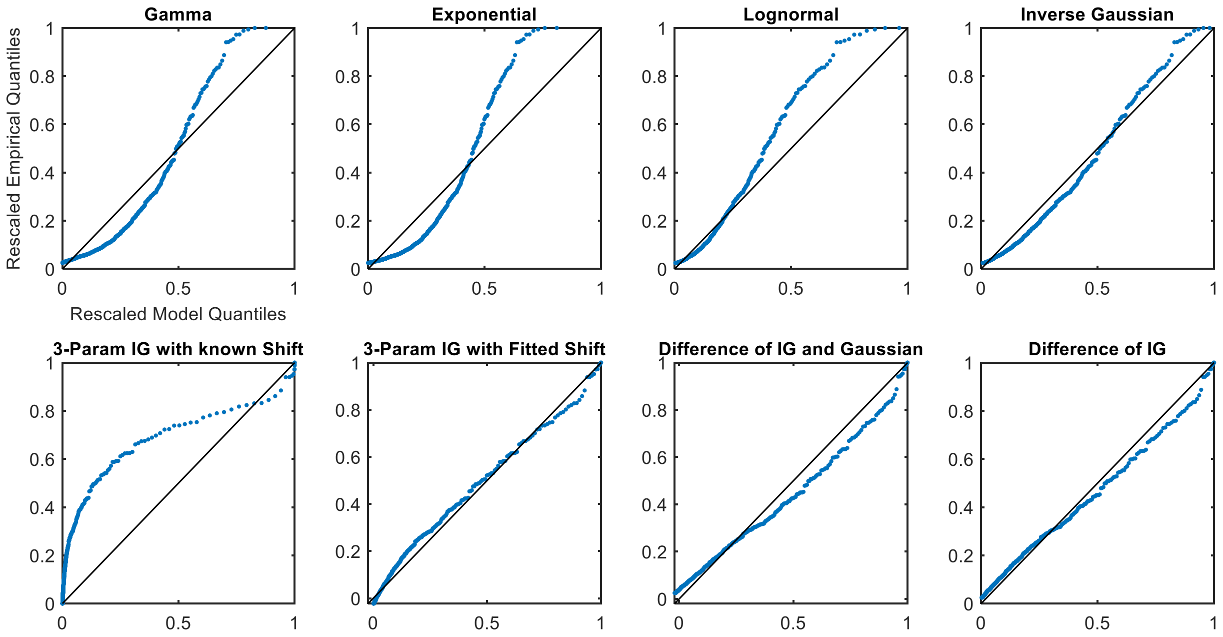


Fig. S28. Rescaled QQ plots for Subject P4 from the propofol sedation cohort

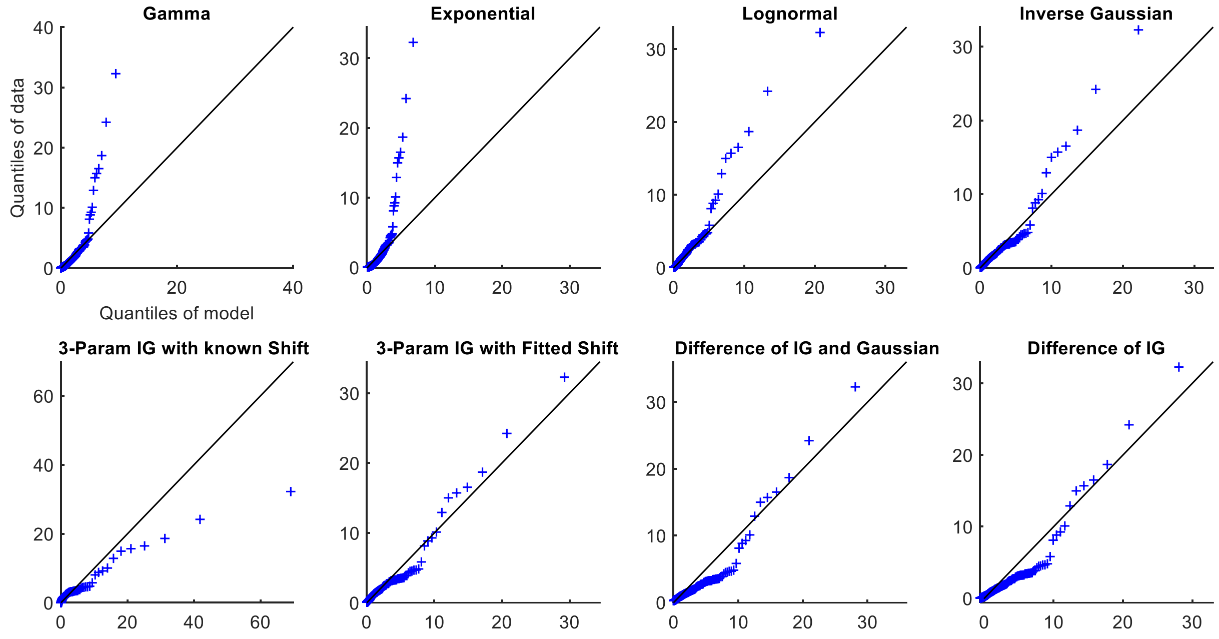


Fig. S29. QQ plots for Subject P5 from the propofol sedation cohort

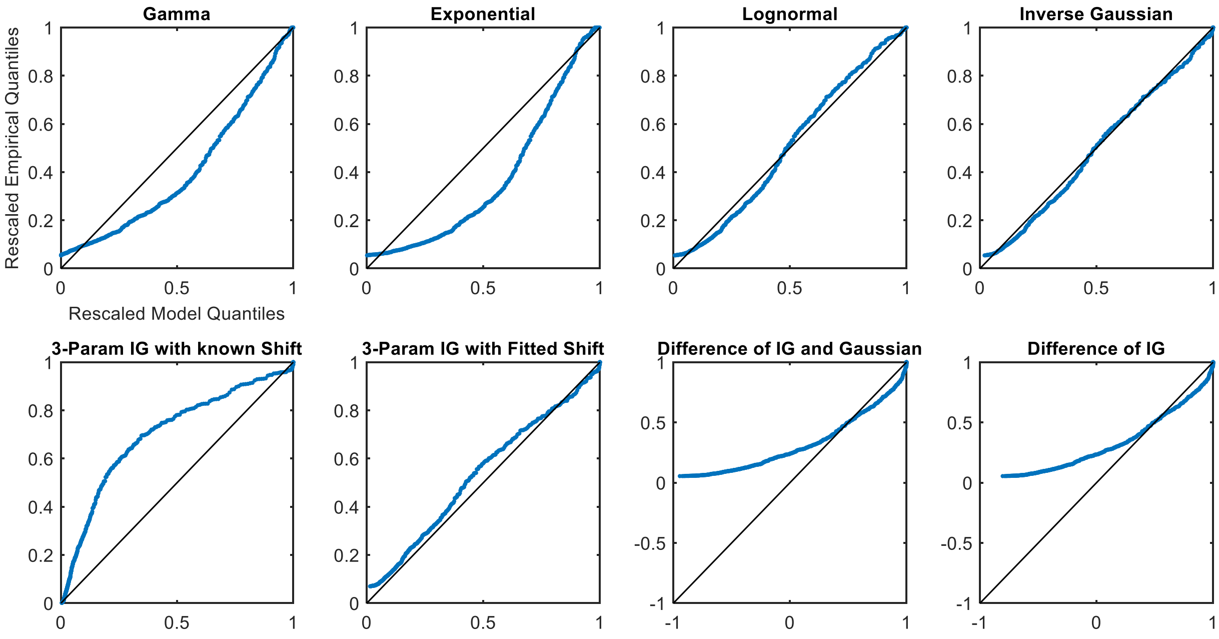


Fig. S30. Rescaled QQ plots for Subject P5 from the propofol sedation cohort

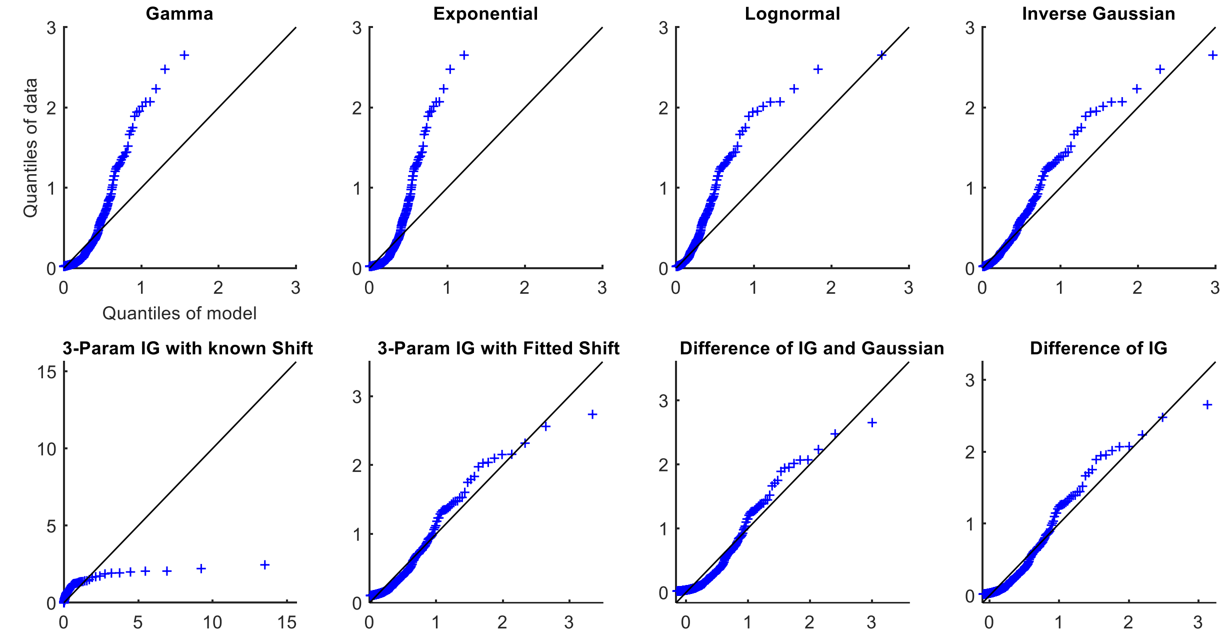


Fig. S31. QQ plots for Subject P6 from the propofol sedation cohort

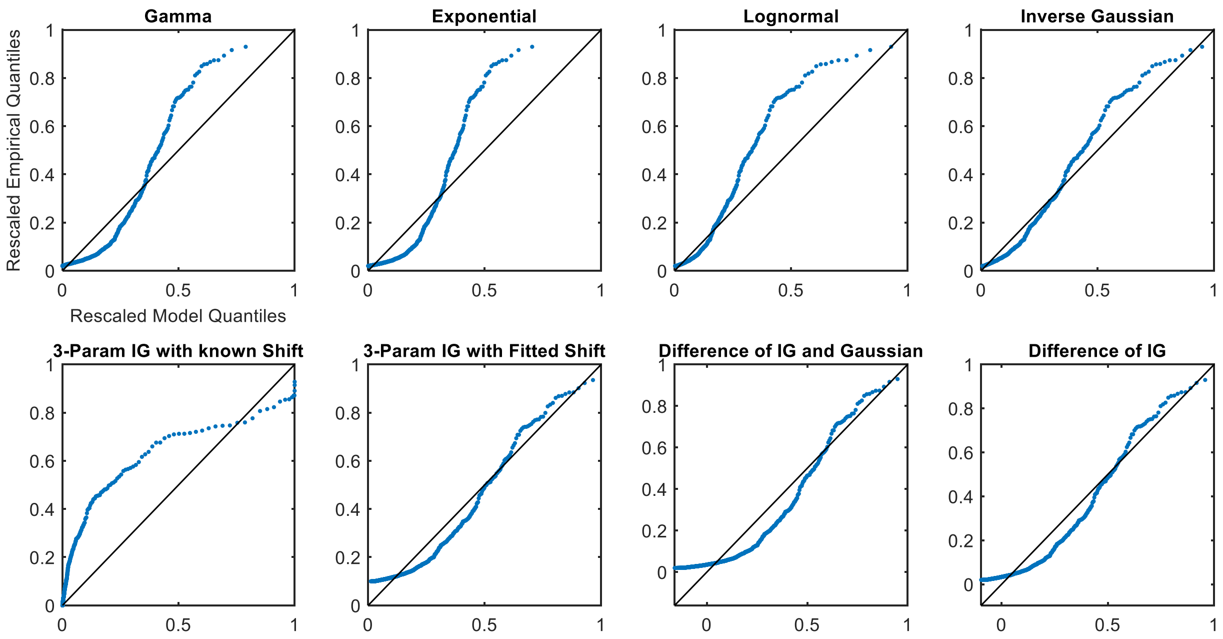


Fig. S32. Rescaled QQ plots for Subject P6 from the propofol sedation cohort

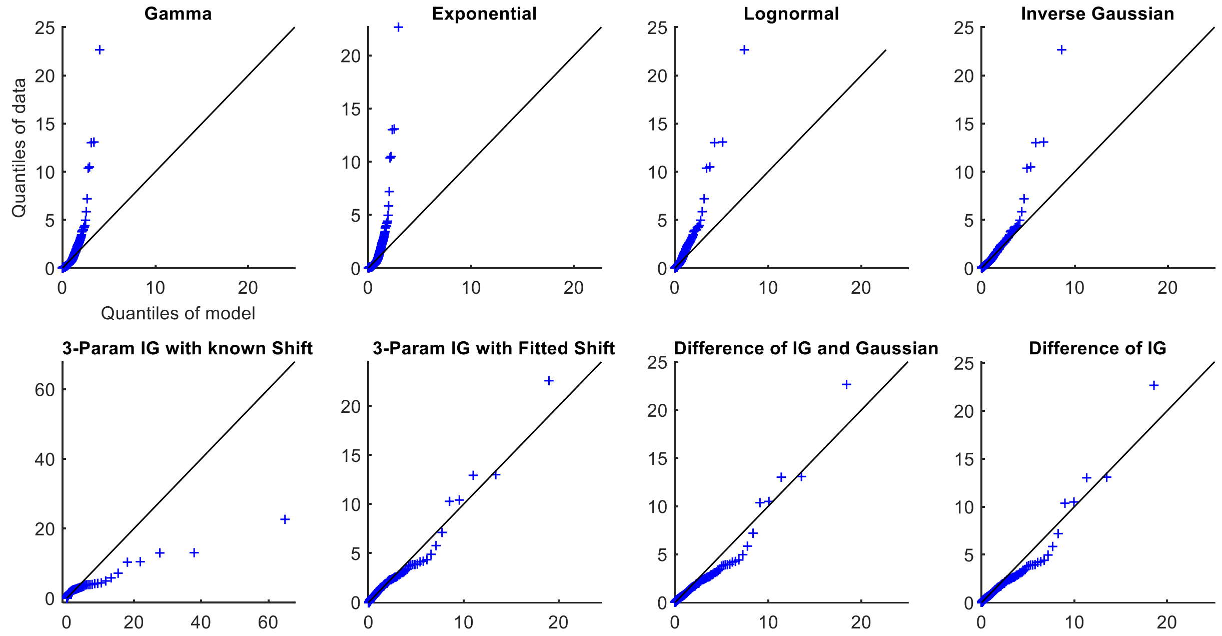


Fig. S33. QQ plots for Subject P7 from the propofol sedation cohort

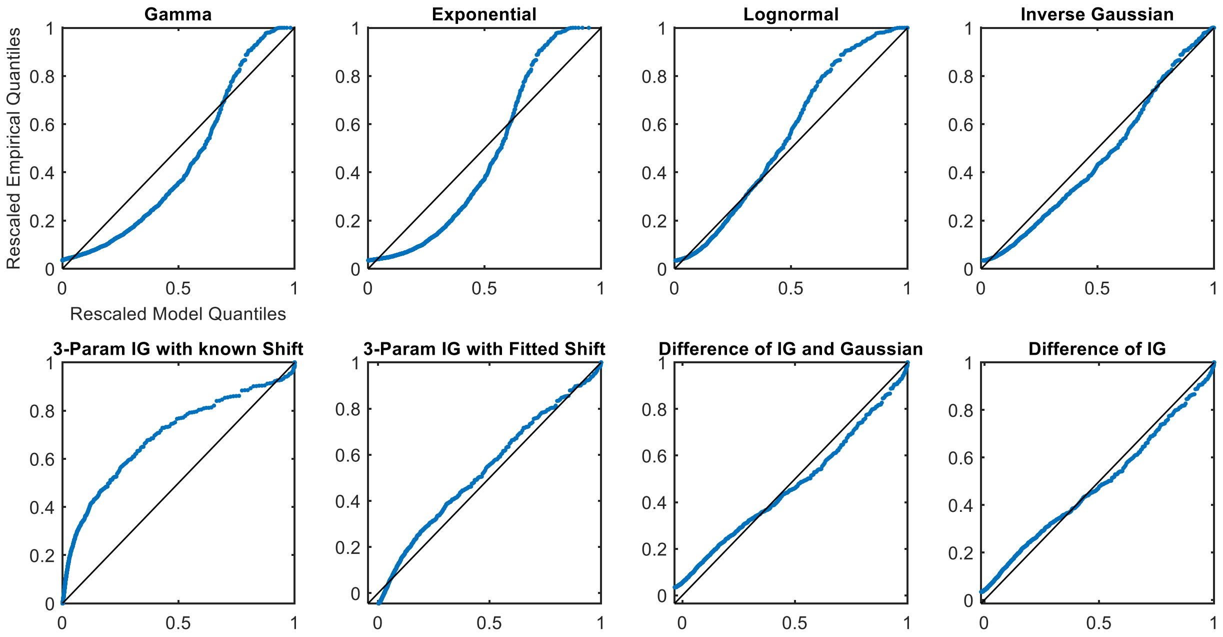


Fig. S34. Rescaled QQ plots for Subject P7 from the propofol sedation cohort

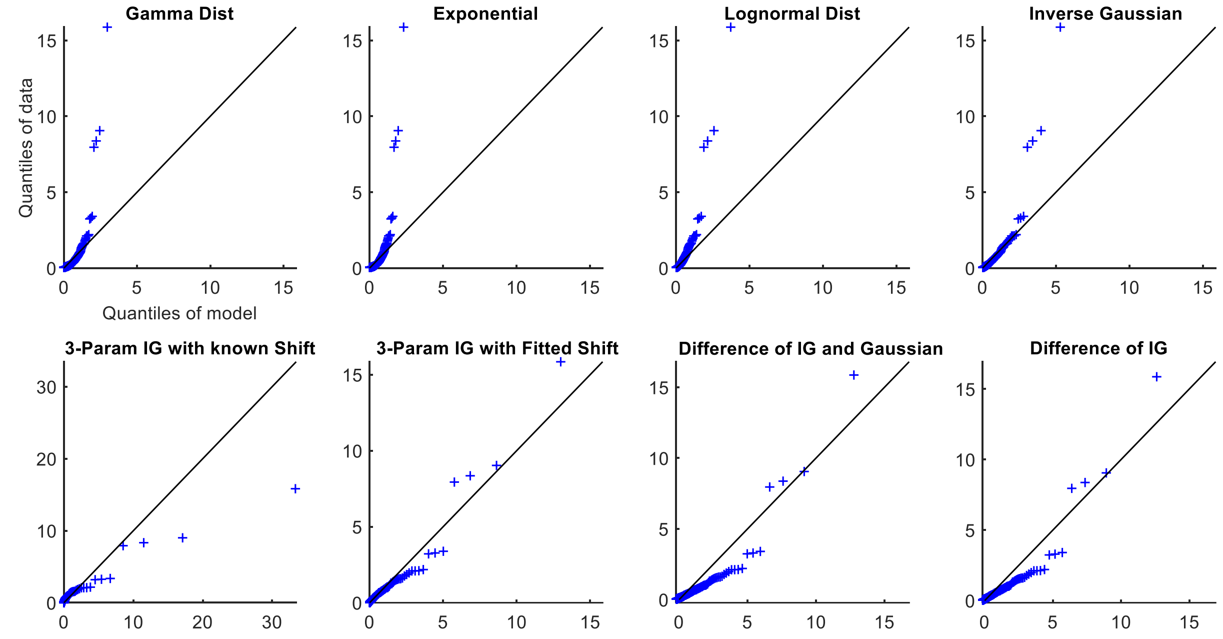


Fig. S35. QQ plots for Subject P8 from the propofol sedation cohort

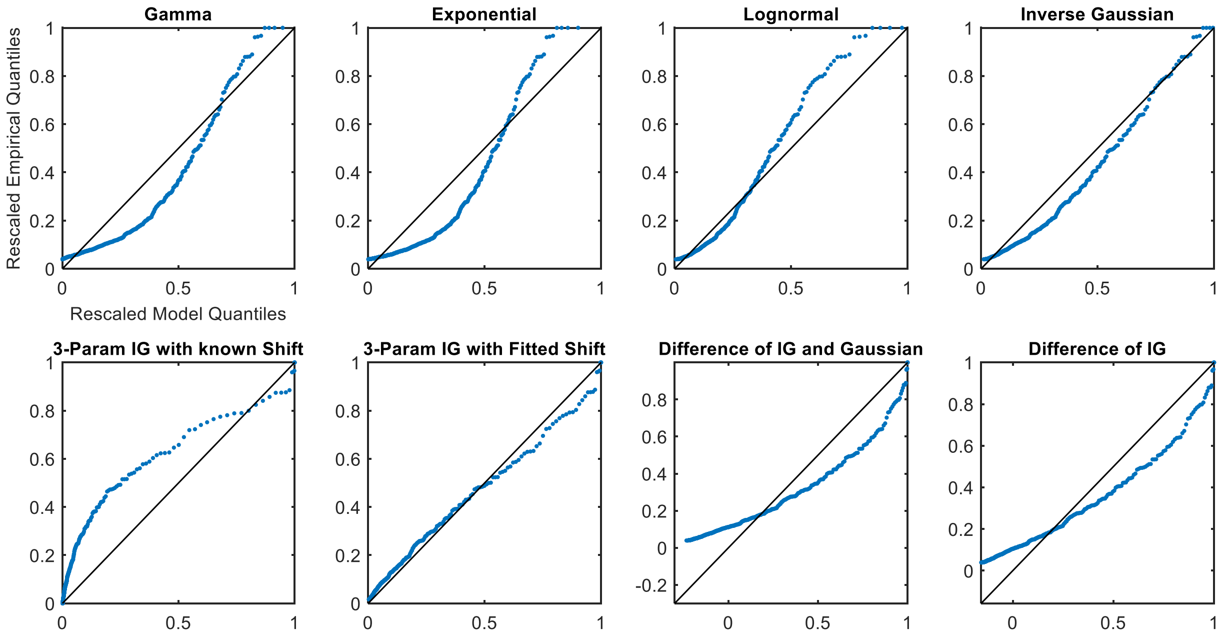


Fig. S36. Rescaled QQ plots for Subject P8 from the propofol sedation cohort

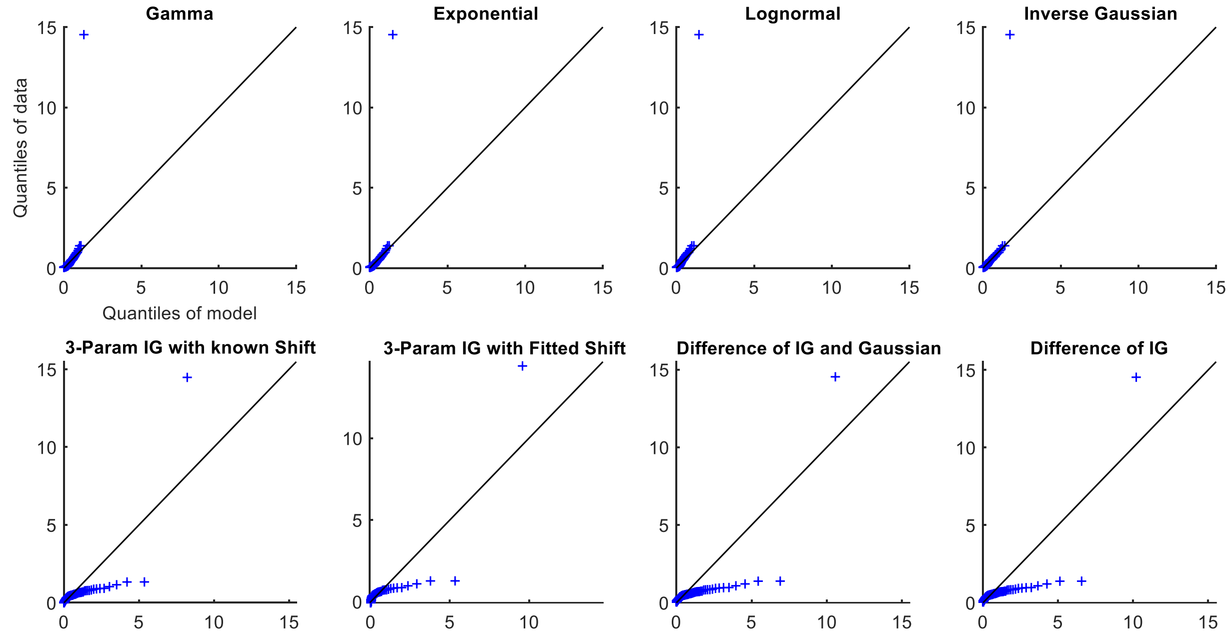


Fig. S37. QQ plots for Subject P9 from the propofol sedation cohort

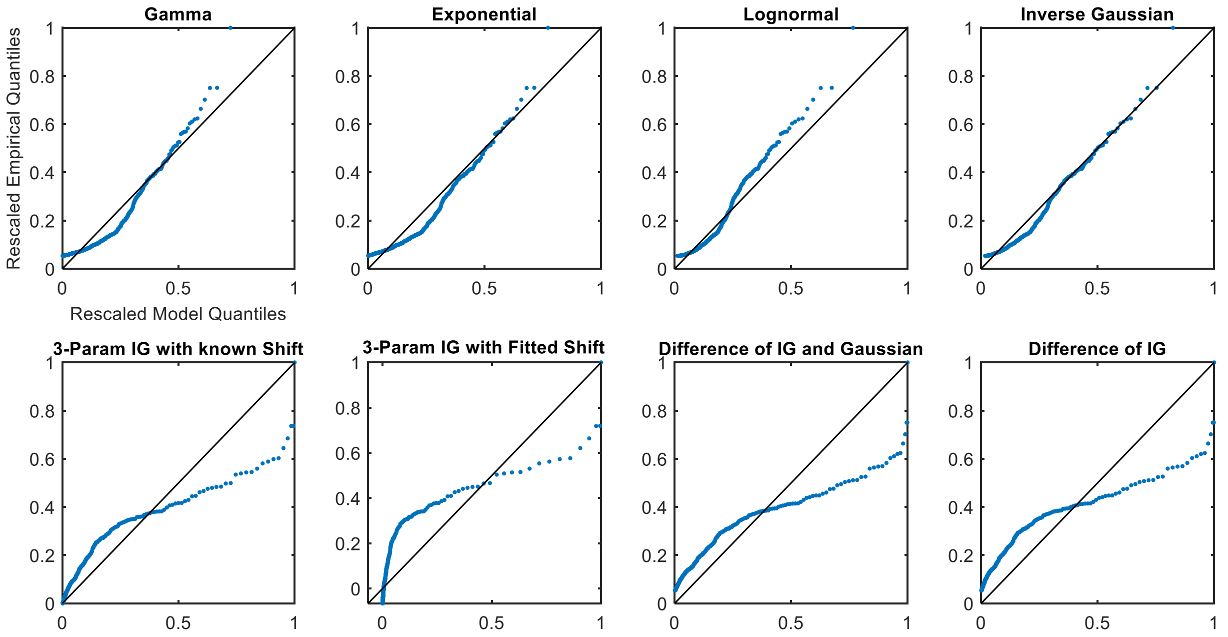


Fig. S38. Rescaled QQ plots for Subject P9 from the propofol sedation cohort

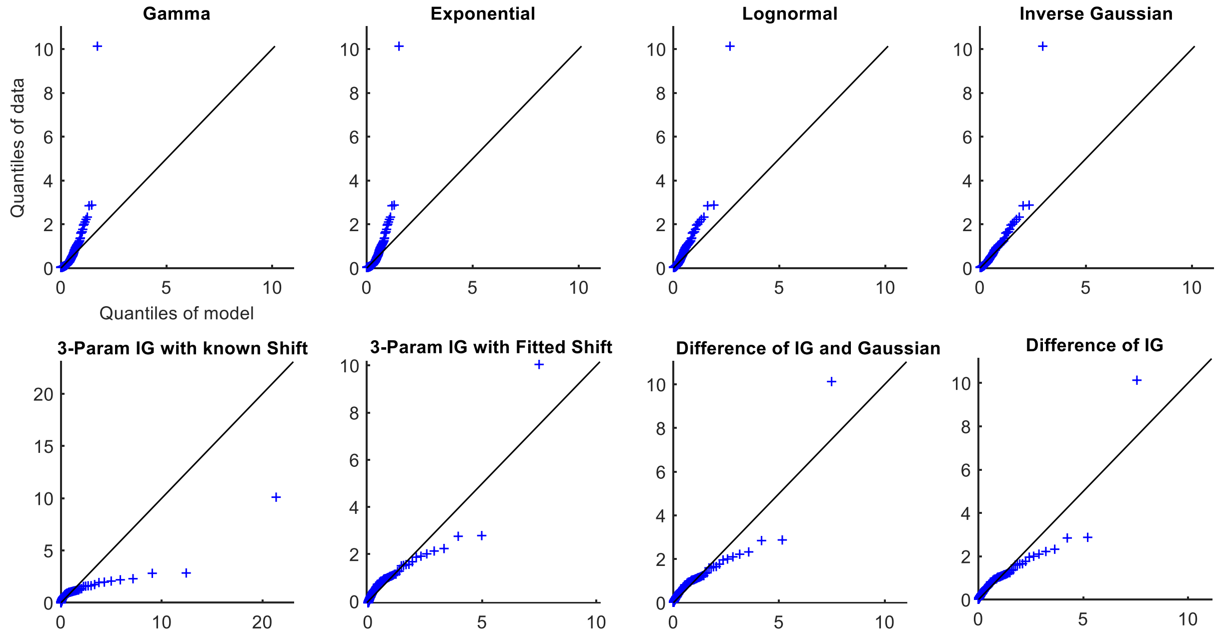


Fig. S39. QQ plots for Subject P11 from the propofol sedation cohort

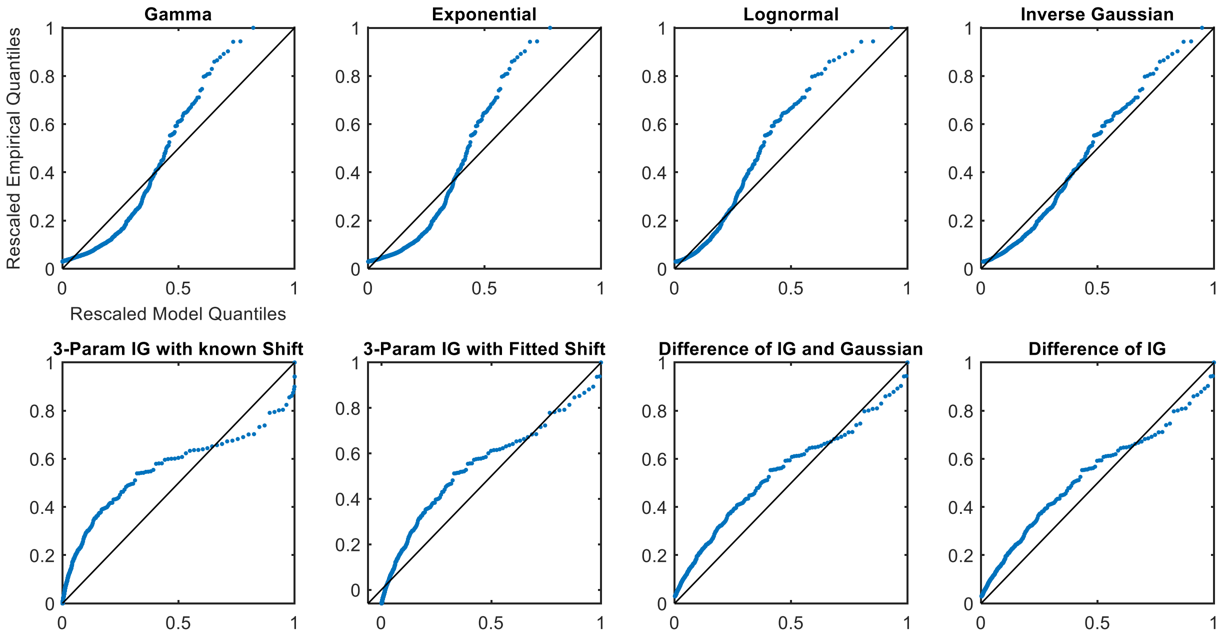


Fig. S40. Rescaled QQ plots for Subject P11 from the propofol sedation cohort
