## Supplementary material for "Elementary Integrate-and-Fire Process Underlies Pulse Amplitudes in Electrodermal Activity": S2 Appendix

Below are the probability densities of all distributions used in the study whose densities can be expressed in closed form.

**Inverse Gaussian**

$$f(x; \mu, \lambda) = \left(\frac{\lambda}{2\pi x^3}\right)^{1/2} \exp\left(-\frac{\lambda(x-\mu)^2}{2\mu^2 x}\right) \qquad \textit{(Eqn. S1)}$$

**3-parameter inverse Gaussian**

$$f(x; \theta, \mu, \lambda) = \left(\frac{\lambda}{2\pi(x-\theta)^3}\right)^{1/2} \exp(-\frac{\lambda[(x-\theta)-\mu]^2}{2\mu^2(x-\theta)}) \qquad \textit{(Eqn. S2)}$$

**Lognormal**

$$f(x; \mu, \sigma) = \left(\frac{1}{2\pi\sigma^2}\right)^{1/2} * \exp\left(-\frac{(\ln x-\mu)^2}{2\sigma^2}\right) * \frac{1}{x} \qquad \textit{(Eqn. S3)}$$

**Gamma**

$$f(x; \alpha, \beta) = \frac{\beta^\alpha}{\Gamma(\alpha)} x^{\alpha-1} e^{-\beta x} \qquad \textit{(Eqn. S4)}$$

**Exponential**

$$f(x; \lambda) = \lambda e^{-\lambda x} \qquad \textit{(Eqn. S5)}$$
