## Supplementary material for "Elementary Integrate-and-Fire Process Underlies Pulse Amplitudes in Electrodermal Activity": S3 Appendix

### 1 Model Formulation

Let and $X$ and $Y$ be inverse Gaussian random variables.

$X \sim IG(\mu_1, \lambda_1)$
$Y \sim IG(\mu_2, \lambda_2)$

Without loss of generality, we define $Z = X - Y$ assuming $\mu_1 > \mu_2$. $Z$ is the difference of two inverse Gaussian distributions and is defined by 4 parameters: $\mu_1, \lambda_1, \mu_2$, and $\lambda_2$. We will now show how to arrive at the method of moments estimates for these 4 parameters.

### 2 Theoretical Moments of the Inverse Gaussian Distribution

Since there are 4 unknowns, we use the first 4 theoretical moments of the inverse Gaussian distribution. These are found in [19] and listed below for $X$ in terms of the moment generating function (MGF) of $X$. We also need the first 4 theoretical moments for $-Y$.

$$
\begin{aligned}
M_X(t) &= \exp\left(\frac{\lambda_1}{\mu_1}\left(1 - \sqrt{1 - \frac{2\mu^2 t}{\lambda_1}}\right)\right) \\
M_X(0) &= 1 \\
M'_X(0) &= \mu_1 \\
M_X^{(2)}(0) &= \mu_1^2 + \frac{\mu_1^3}{\lambda_1} \\
M_X^{(3)}(0) &= \mu_1^3 + \frac{3\mu_1^4}{\lambda_1} + \frac{3\mu_1^5}{\lambda_1^2} \\
M_X^{(4)}(0) &= \mu_1^4 + \frac{6\mu_1^5}{\lambda_1} + \frac{15\mu_1^6}{\lambda_1^2} + \frac{15\mu_1^7}{\lambda_1^3}
\end{aligned}
$$

$$
\begin{aligned}
M_{-Y}(0) &= M_Y(0) * (-1)^0 = 1 \\
M'_{-Y}(0) &= M'_Y(0) * (-1)^1 \\
&= -\mu_2 \\
M_{-Y}^{(2)}(0) &= M_Y^{(2)}(0) * (-1)^2 \\
&= \mu_2^2 + \frac{\mu_2^3}{\lambda_2} \\
M_{-Y}^{(3)}(0) &= M_Y^{(3)}(0) * (-1)^3 \\
&= -\mu_2^3 - \frac{3\mu_2^4}{\lambda_2} - \frac{3\mu_2^5}{\lambda_2^2} \\
M_{-Y}^{(4)}(0) &= M_Y^{(4)}(0) * (-1)^4 \\
&= \mu_2^4 + \frac{6\mu_2^5}{\lambda_2} + \frac{15\mu_2^6}{\lambda_2^2} + \frac{15\mu_2^7}{\lambda_2^3}
\end{aligned}
$$

#### 3 Deriving the Theoretical Moments of the Difference Distribution

We know that the MGF of $Z$ is the product of the MGF's of $X$ and $-Y$.

$M_Z(t) = M_X(t) * M_{-Y}(t)$

Using this, we can compute the first four theoretical moments of $Z$.

$$
\begin{aligned}
M'_Z(0) &= M'_X(0)M_{-Y}(0) + M_X(0)M'_{-Y}(0) \\
&= \mu_1 * 1 + 1 * -\mu_2 \\
&= \mu_1 - \mu_2 \\
M_Z^{(2)}(0) &= M_X^{(2)}(0)M_{-Y}(0) + 2M'_X(0)M'_{-Y}(0) + M_X(0)M_{-Y}^{(2)}(0) \\
&= (\mu_1^2 + \frac{\mu_1^3}{\lambda_1})(1) + 2(\mu_1)(-\mu_2) + (1)(\mu_2^2 + \frac{\mu_2^3}{\lambda_2}) \\
&= (\mu_1 - \mu_2)^2 + \frac{\mu_1^3}{\lambda_1} + \frac{\mu_2^3}{\lambda_2} \\
M_Z^{(3)}(0) &= M_X^{(3)}(0)M_{-Y}(0) + 3M_X^{(2)}(0)M'_{-Y}(0) + 3M'_X(0)M_{-Y}^{(2)}(0) + M_X(0)M_{-Y}^{(3)}(0) \\
&= (\mu_1^3 + \frac{3\mu_1^4}{\lambda_1} + \frac{3\mu_1^5}{\lambda_1^2})(1) + 3(\mu_1^2 + \frac{\mu_1^3}{\lambda_1})(-\mu_2) + 3(\mu_1)(\mu_2^2 + \frac{\mu_2^3}{\lambda_2}) + (1)(-\mu_2^3 - \frac{3\mu_2^4}{\lambda_2} - \frac{3\mu_2^5}{\lambda_2^2}) \\
&= (\mu_1 - \mu_2)^3 + 3(\frac{\mu_1^3}{\lambda_1} + \frac{\mu_2^3}{\lambda_2})(\mu_1 - \mu_2) + 3(\frac{\mu_1^5}{\lambda_1^2} - \frac{\mu_2^5}{\lambda_2^2}) \\
M_Z^{(4)}(0) &= M_X^{(4)}(0)M_{-Y}(0) + 4M_X^{(3)}(0)M'_{-Y}(0) + 6M_X^{(2)}(0)M_{-Y}^{(2)}(0) + 4M'_X(0)M_{-Y}^{(3)}(0) + M_X(0)M_{-Y}^{(4)}(0) \\
&= (\mu_1 - \mu_2)^4 + 15(\frac{\mu_1^7}{\lambda_1^3} + \frac{\mu_2^7}{\lambda_2^3}) + 6(\frac{\mu_1^3}{\lambda_1} + \frac{\mu_2^3}{\lambda_2})(\mu_1 - \mu_2)^2 + 3(\frac{\mu_1^3}{\lambda_1} + \frac{\mu_2^3}{\lambda_2})^2 + 12(\mu_1 - \mu_2)(\frac{\mu_1^5}{\lambda_1^2} - \frac{\mu_2^5}{\lambda_2^2})
\end{aligned}
$$

#### 4 Method of Moments for the Difference of Two Inverse Gaussians

Setting the theoretical moments computed above equal to the respective empirical moments ($x_1, x_2, ..., x_n$ are the data points), we arrive at the following system of 4 equations with 4 unknowns:

Letting $\bar{x} = \Sigma_{i=1}^n x_i/n$,

$$
\begin{aligned}
\mu_1 - \mu_2 &= \bar{x} \\
(\mu_1 - \mu_2)^2 + \frac{\mu_1^3}{\lambda_1} + \frac{\mu_2^3}{\lambda_2} &= \frac{\Sigma_{i=1}^n x_i^2}{n} \\
(\mu_1 - \mu_2)^3 + 3(\frac{\mu_1^3}{\lambda_1} + \frac{\mu_2^3}{\lambda_2})(\mu_1 - \mu_2) + 3(\frac{\mu_1^5}{\lambda_1^2} - \frac{\mu_2^5}{\lambda_2^2}) &= \frac{\Sigma_{i=1}^n x_i^3}{n} \\
(\mu_1 - \mu_2)^4 + 15(\frac{\mu_1^7}{\lambda_1^3} + \frac{\mu_2^7}{\lambda_2^3}) + 6(\frac{\mu_1^3}{\lambda_1} + \frac{\mu_2^3}{\lambda_2})(\mu_1 - \mu_2)^2 + 3(\frac{\mu_1^3}{\lambda_1} + \frac{\mu_2^3}{\lambda_2})^2 + 12(\mu_1 - \mu_2)(\frac{\mu_1^5}{\lambda_1^2} - \frac{\mu_2^5}{\lambda_2^2}) &= \frac{\Sigma_{i=1}^n x_i^4}{n}
\end{aligned}
$$

After much simplification and substitution, we arrive at the following more manageable system of 4 equations with 4 unknowns:

$$
\begin{aligned}
\mu_1 - \mu_2 &= \bar{x} \\
\frac{\mu_1^3}{\lambda_1} + \frac{\mu_2^3}{\lambda_2} &= \frac{\Sigma_{i=1}^n x_i^2}{n} - \bar{x}^2 \\
3(\frac{\mu_1^5}{\lambda_1^2} - \frac{\mu_2^5}{\lambda_2^2}) &= (\frac{\Sigma_{i=1}^n x_i^3}{n} - \bar{x}^3) - 3\bar{x}(\frac{\Sigma_{i=1}^n x_i^2}{n} - \bar{x}^2) \\
15(\frac{\mu_1^7}{\lambda_1^3} + \frac{\mu_2^7}{\lambda_2^3}) &= (\frac{\Sigma_{i=1}^n x_i^4}{n} - \bar{x}^4) - 4\bar{x}(\frac{\Sigma_{i=1}^n x_i^3}{n} - \bar{x}^3) + 6\bar{x}^2(\frac{\Sigma_{i=1}^n x_i^2}{n} - \bar{x}^2) - 3(\frac{\Sigma_{i=1}^n x_i^2}{n} - \bar{x}^2)^2
\end{aligned}
$$

While not closed form, solving this system yields the method of moments estimates of the 4 parameters of the difference of two inverse Gaussians. Since all parameters must be positive, this system can be solved with a numerical least squares system solver in Matlab or Python.

#### 5 Gaussian Approximation for the Second Inverse Gaussian

Now, let's say $\tilde{Y} \sim N(\mu_2, \sigma_2^2)$, and we want to compute the method of moments estimates for $\tilde{Z} = X - \tilde{Y}$. We know that $M_{\tilde{Z}}(t) = M_X(t) * M_{-\tilde{Y}}(t)$. The first 4 theoretical moments of $-\tilde{Y}$ are below.

$$
\begin{aligned}
M_{\tilde{Y}}(t) &= \exp{(\mu_2 t + \frac{\sigma_2^2 t^2}{2})} \\
M_{-\tilde{Y}}(0) &= M_{\tilde{Y}}(0) * (-1)^0 = 1 \\
M'_{-\tilde{Y}}(0) &= M'_{\tilde{Y}}(0) * (-1)^1 \\
&= -\mu_2 \\
M^{(2)}_{-\tilde{Y}}(0) &= M^{(2)}_{\tilde{Y}}(0) * (-1)^2 \\
&= \mu_2^2 + \sigma_2^2 \\
M^{(3)}_{-\tilde{Y}}(0) &= M^{(3)}_{\tilde{Y}}(0) * (-1)^3 \\
&= -\mu_2^3 - 3\mu_2\sigma_2^2 \\
M^{(4)}_{-\tilde{Y}}(0) &= M^{(4)}_{\tilde{Y}}(0) * (-1)^4 \\
&= \mu_2^4 + 6\mu_2^2\sigma_2^2 + 3\sigma_2^4
\end{aligned}
$$

The first 4 theoretical moments of $\tilde{Z}$ are below.

$$M'_Z(0) = \mu_1 - \mu_2$$
$$M_Z^{(2)}(0) = (\mu_1 - \mu_2)^2 + \frac{\mu_1^3}{\lambda_1} + \sigma_2^2$$
$$M_Z^{(3)}(0) = (\mu_1 - \mu_2)^3 + 3(\mu_1 - \mu_2)(\frac{\mu_1^3}{\lambda_1} + \sigma_2^2) + 3\frac{\mu_1^5}{\lambda_1^2}$$
$$M_Z^{(4)}(0) = (\mu_1 - \mu_2)^4 + 15\frac{\mu_1^7}{\lambda_1^3} + 6(\frac{\mu_1^3}{\lambda_1} + \sigma_2^2)(\mu_1 - \mu_2)^2 + 3(\frac{\mu_1^3}{\lambda_1} + \sigma_2^2)^2 + 12\frac{\mu_1^5}{\lambda_1^2}(\mu_1 - \mu_2)$$

Setting the first 4 theoretical moments of $\tilde{Z}$ to the respective empirical moments, we arrive at the following system of 4 equations with 4 unknowns.

$$\mu_1 - \mu_2 = \bar{x}$$
$$\frac{\mu_1^3}{\lambda_1} + \sigma_2^2 = \frac{\Sigma_{i=1}^n x_i^2}{n} - \bar{x}^2$$
$$3\frac{\mu_1^5}{\lambda_1^2} = (\frac{\Sigma_{i=1}^n x_i^3}{n} - \bar{x}^3) - 3\bar{x}(\frac{\Sigma_{i=1}^n x_i^2}{n} - \bar{x}^2)$$
$$15\frac{\mu_1^7}{\lambda_1^3} = (\frac{\Sigma_{i=1}^n x_i^4}{n} - \bar{x}^4) - 4\bar{x}(\frac{\Sigma_{i=1}^n x_i^3}{n} - \bar{x}^3) + 6\bar{x}^2(\frac{\Sigma_{i=1}^n x_i^2}{n} - \bar{x}^2) - 3(\frac{\Sigma_{i=1}^n x_i^2}{n} - \bar{x}^2)^2$$
